# Anterior thalamic nucleus inputs to the retrosplenial cortex are unperturbed in the hAPP-J20 mouse model of Alzheimer’s disease

**DOI:** 10.64898/2025.12.01.691675

**Authors:** Gabriella Margetts-Smith, Lilya Andrianova, Shivali Kohli, Andrew D Randall, John P Aggleton, Jonathan Witton, Michael T Craig

## Abstract

Alzheimer’s disease (AD) is a neurodegenerative condition characterised by progressive loss of memory and general decline in cognitive function. Although research has traditionally focused on the hippocampus and entorhinal cortex, other regions important for memory and spatial navigation, such as the anterior thalamic nuclei (ATN) and retrosplenial cortex (RSC), are also affected. The RSC is an important brain region for both memory and spatial navigation, and displays dysfunction in humans during prodromal phases of AD. The ATN provide a major input to RSC, with loss of this input to RSC causing a dramatic reduction in the expression of the immediate early gene product c-Fos. The ATN to RSC connection may thus present a new target for therapeutic intervention in AD.

Here, we set out to determine whether AD-like pathology perturbed projections from ATN to RSC using the hAPP-J20 amyloidopathy mouse model, using optogenetic activation of thalamic inputs combined with *ex vivo* patch clamp electrophysiology. We found that RSC displayed an age-dependent decline in basal c-Fos activity, with no additional effect of amyloid pathology on c-Fos expression. At all time points (3, 6 and 9 months) measured, we found no evidence of impairments in the synaptic strength or efficacy of the ATN projections to either granular or dysgranular subdivisions of RSC. We conclude that, in the hAPP-J20 mouse model, this pathway is unaffected by amyloid-β overexpression.

## Introduction

Alzheimer’s disease (AD) is a progressive neurodegenerative disease that typically initiates in entorhinal and hippocampal regions^1^ and also specifically affects the midline and anterior thalamic nuclei that project to these regions^2,3^. Impairments in memory are often the first symptoms that present in individuals living with Alzheimer’s disease^4^, although the consensus is that pathology may be present for 10 to 20 years before the onset of symptoms^5^. Understanding how AD affects the neural circuitry underlying memory at different points in disease progression is an essential first step to developing targeted, circuits-based treatments to allow amelioration of memory deficits once the disease has begun to progress. While research into the neurobiology of memory has typically focused on the hippocampus, there are numerous other, equally important brain regions such as the entorhinal and retrosplenial cortices, that are involved in the formation and maintenance of memory. Alongside the midline thalamus, the anterior thalamic nuclei (ATN) play a key role in serving as a hub for communication throughout the brain’s extended memory and present an attractive target for therapeutic intervention into spatial and episodic memory disorders^6^.

One brain region of particular interest in the extended memory circuit is the retrosplenial cortex (RSC). The retrosplenial cortex is highly interconnected with hippocampal, parahippocampal and anterior thalamic regions, with this connectivity highly conserved across mammalian species^7^. The RSC also receives input from sensory regions such as visual cortex and is believed to have important roles in both memory and spatial navigation^7^. In humans, the RSC is site of reduced metabolism in the early stages of AD^8^, with impaired functional connectivity sufficient to distinguish RSC from posterior cingulate cortex in prodromal AD^9^. Other studies have found that prodromal AD is associated with RSC hypometabolism, atrophy, synapse loss and Aβ accumulation^10–13^. A recent combined human and rodent study found dysregulation of retrosplenial cortex in AD was associated with impairments in the function of inhibitory GABAergic neurons^14^. This latter study highlights the need for animal studies to better understand the function of RSC in health and disease, given the difficulty of directly manipulating RSC in humans^3,7^.

In mouse models of amyloidopathy, the RSC exhibits significant Aβ deposition which is concurrent with neuroinflammatory responses as well as cognitive impairment in the 5xFAD model^15^. The RSC also displays decreased neuronal activity in response to a novel environment at a preclinical stage preceding plaque deposition in the Tg2576 amyloidopathy model^16^. Indeed, the RSC is sensitive to loss of synaptic inputs, with deafferentation via hippocampal^17^ or anterior thalamic nuclear lesions leading to a dramatic reduction in the expression of the immediate-early gene product c-Fos.

Altogether, it is clear that retrosplenial cortex may be important both in pathophysiology of Alzheimer’s disease as well as presenting a tempting therapeutic target. In the present study, we used the hAPP-J20 amyloidogenic mouse model^18^ of AD to determine whether markers of activity in RSC were impaired at time points relating to pre-symptomatic, prodromal and symptomatic phases of the disorder. We also used optogenetic activation of ATN inputs to the RSC, combined with patch-clamp electrophysiology, to test the hypothesis that the ATN to RSC projection would specifically be impaired in this mouse model.

## Methods

### Animals

All experiments were conducted in accordance with the UK Animals (Scientific Procedures) Act 1986 under Project licence PAC082CD3, after local ethical review by the Animal Welfare and approved by the University of Exeter Institutional Ethical Review Board. All animals were maintained on a 12h constant light/dark cycle and procedures and experiments were conducted during the light phase. All animals had access to food and water *ad libitum* and were group-housed wherever possible. We used standard home-cage enrichment that included cardboard tubes and domes, wooden chew blocks and nesting material.

For immunohistochemistry experiments, we used 49 wild-type (WT) and heterozygous transgenic hAPP-J20^18^ mice. Founder mice for our hAPP-J20 colony were kindly supplied by Dr Keith Phillips of Eli Lilly (Erl Wood, Surrey, UK) on a C57BL/6J background and were occasionally back-crossed with C57BL/6J mice to maintain the background. We used heterozygous and WT littermates that were bred from wildtype X heterozygote pairings.

### Drugs and Chemicals

CGP55845, Gabazine (SR 95531) and DNQX were purchased from HelloBio (UK), and all other chemicals were purchased from Sigma-Aldrich (UK) unless otherwise stated.

### Immunohistochemistry

For immunohistochemistry experiments, we used 49 wild-type (WT) and hAPP-J20 mice at 3 months (3m), 6 months (6m) and 9 months (9m). 24 WT and 25 hAPP-J20 mice were deeply anaesthetised with sodium pentobarbitone (200 mg/kg; Henry Schein) before being transcardially perfused (5 ml/min) with ice-cold phosphate buffered saline (PBS) for 5 mins followed by 4% paraformaldehyde (PFA) in PBS for 10 mins. The animals were not handled or exposed to any other non-home cage stimulation for 2 hours prior to perfusion in order to capture only basal c-Fos expression. For the c-Fos experiment, mice were group-housed (2 to 5 mice per cage) in GM500 individually vented cages (Techniplast, UK) with dimensions of 399 x 199 x 169 mm. The cages had standard enrichment of cardboard tubes, mouse shelter and bedding. Following the perfusion, the brains were removed and stored in 4% PFA in PBS for 22 hours at 4°C, then cryoprotected in 30% Sucrose (in PBS with0.02% Sodium Azide for at least 3 days. Mice were genotyped prior to perfusion as part of general animal husbandry practices, and genotype was confirmed from tissue removed from the tail after anaesthesia was applied. 30 µm coronal sections were collected and stained for Fos and Aβ plaques, and tissue and staining quality was assessed from Fos-stained sections. Mice were excluded from analysis of Fos and Aβ deposition if sections were highly damaged or if the Fos staining failed (assessed from an absence of staining in the olfactory bulb, piriform cortex, somatosensory cortex and visual cortex). 9 mice were excluded due to low tissue quality, leaving 18 WT (3m: n_male_ = 4, n_female_ = 5; 6m: n_male_ = 1, n_female_ = 3; 9m: n_male_ = 4, n_female_ = 1) and 22 Tg mice (3m: n_male_ = 2, n_female_ = 5; 6m: n_male_ = 4, n_female_ = 3; 9m: n_male_ = 5, n_female_ = 3).

Using a freezing sledge microtome (Leica SM2010R with Physitemp BFS-5MP temperature controller) coronal sections were taken from frozen brains (-20°C). Free-floating sections were stored in PBS containing (0.02%) NaAzide at 4°C or a cryoprotectant solution (25% glycerol, 30% ethylene glycol, 25% 0.2 M Phosphate buffer, 20% ddH_2_O) at -20°C for long-term storage. For immunohistochemistry staining, all steps were conducted at RT unless otherwise stated.

For c-Fos staining, 30 µm free-floating sections stored in PBS-Azide were washed in PBS (3x 10 minutes), before being incubated in PBS containing 0.09% hydrogen peroxide for 20 minutes to quench endogenous peroxidase. Next, sections were washed in PBS (3x 10 minutes), then blocked and permeabilised in PBS-(0.2%)Tx (PBS, 0.2% Triton X-100) containing 3% normal goat serum (NGS). The sections were then incubated in 1:800 anti-Fos primary antibody in PBS-(0.2%)Tx with 3% NGS at 4°C overnight. The following day the sections were washed in PBS (3x 10 minutes), then incubated for 2 hours in 1:600 biotinylated anti-rabbit secondary antibody in PBS-(0.2%)Tx with 1% NGS. Sections were washed in PBS (2x 10 minutes), then incubated in an Avidin-Biotin complex (ABC) solution for 1 hour. After a further three washes in PBS the sections were incubated with 0.04% 3,3’-Diaminobenzidine-tetrahydrochloride (DAB) with 0.04% hydrogen peroxide and 0.05% ammonium nickel(II) sulfate in PBS for approximately 10 minutes. After a final two 10-minute washes in PBS, sections were mounted on Superfrost Plus slides and left to dry overnight. The next day, sections were serially dehydrated in graded ethanol (EtOH) baths (all 2 minutes: 2x ddH_2_O, 30% EtOH, 60% EtOH, 90% EtOH, 95% EtOH, 2x 100% EtOH) and then cleared in Histo-Clear II (2x 10 minutes). Slides were then sealed and coverslipped using Histo-Mount mounting medium.

The RSC, CA1 region of the hippocampus (HPC) and the entorhinal cortex (EC) were visualised using bright-field microscopy and images captured using a 10x objective on a Nikon Eclipse 800 microscope attached to a SPOT RT monochrome camera running SPOT Basic imaging capture software (SPOT Imaging). Four replicates of each ROI were captured for each mouse, taken from the left and right hemispheres of two sections. Image analysis consisted of an automatic count of nuclei expressing high levels of c-Fos (Fos+) in predefined regions of interest (ROI) using Paxinos & Franklin boundaries^19^ on Fiji^20^ software. To perform this count, images were submitted to a fast Fourier transform bandpass filter and colour scale inversion before being run through the 3D object counter plugin with a brightness threshold (set to 1.7) that depended on average pixel brightness of the filtered image^21^. Fos^+^ cell counts were then normalised to ROI area, described as Fos^+/mm2^.

For amyloid plaque staining, 30 µm free-floating sections stored in cryoprotectant solution were washed in PBS (3x 10 minutes) before being mounted onto Superfrost Plus slides and left to dry overnight. The following morning a hydrophobic barrier was drawn around the sections and sections were covered in 70% ethanol for 5 minutes before being washed with ddH_2_O (2x 2 minutes). Next, sections were incubated in 1:100 Amylo-Glo^22^ (2B Scientific) solution in 0.9% saline for 10 minutes. Sections were then rinsed in 0.9% saline for 5 minutes, followed by PBS washes (3x 10 minutes). Sections were left to dry, then cover slipped with Fluoromount mounting medium (Merck Life Sciences).

Aβ plaques in the RSC, CA1 and EC were visualised using epifluorescent microscopy (Nikon Eclipse 800 microscope with CoolLED pE-4000 LED light source) at 365 nm excitation. Images were captured at 4x magnification using a SPOT RT camera running SPOT Basic imaging capture software. Four replicates of each ROI were captured for each mouse. Image analysis was conducted using Fiji software; amyloid plaques were quantified by measuring the area covered by plaques in predefined brain ROIs. Plaque areas were manually drawn and measured where present, then normalised as a percentage of brain area, described as Plaque/Area (%).

### Stereotaxic injection surgical procedure

All surgeries were conducted using aseptic technique. Mice were anaesthetised with isoflurane (5% induction, 1.2-2.5% maintenance) delivered in a constant flow of oxygen. The mice were placed on a heated pad (37°C) for the duration of the surgery and given 0.03 mg/kg of buprenorphine (buprenorphine hydrochloride, Henry Schein) subcutaneously at the beginning of surgery as an adjunct analgesic, plus 5 mg/kg of carprofen (Rimadyl, Henry Schein) and 10 ml/kg of 0.9% saline were given subcutaneously immediately post-surgery. 5 mg/kg of carprofen was also provided daily for 3 days post-surgery.

Mice received a unilateral injection into the ATN (AD/AV border; AP: -0.71 mm; ML: ±0.80 mm; DV -2.76 mm from pia) of 250 nl of AAV_5_-hSyn1-ChR2(H134R)-mCherry-WPRE-hGHp(A) (ETH Zurich Viral Vector Facility, titre 7.1 x 10^12^ vg/ml) at a flow rate of 50 nl/min. After the surgery, the mice were allowed at a 4-7 week recovery period to allow sufficient time for the expression of the viral construct, with electrophysiological recordings carried out in mice aged 3, 6, or 9 months.

### Slice preparation and electrophysiology

Mice were anaesthetised with 5% isoflurane and decapitated before the brain was rapidly removed and placed in room temperature (RT) oxygenated N-methyl-d-glucamine (NMDG) solution^23^ consisting of (in mM): 135 NMDG, 10 D-Glucose, 1.5 MgCl_2_, 0.5 CaCl_2_, 1 KCl, 1.2 KH_2_PO_4_, 20 Choline bicarbonate. 300 µm coronal slices corresponding to approximately Bregma -0.5 mm to -4.5 mm were sectioned using a Leica VT1200 vibratome. Following sectioning, slices were transferred to a holding chamber containing artificial cerebral spinal fluid (aCSF) perfused with a continuous flow of carbogen (95% O_2_, 5% CO_2_). The aCSF solution consisted of (in mM): 119 NaCl, 3 KCl, 1 NaH_2_PO_4_, 26 NaHCO_3_, 10 D-Glucose, 2.5 CaCl_2_, 1.3 MgCl (pH 7.4). The slices were then incubated at 35°C for 30 minutes, then at RT for a further 30 minutes to recover before recording. ATN injection site fluorescence was visually confirmed immediately following slicing, and slices were discarded if viral expression was absent or in an anatomically-incorrect location (n=24 mice). Slices from a total of 68 mice were used: 39 WT (3m: n_male_ = 8, n_female_ = 3; 6m: n_male_ = 7, n_female_ = 9; 9m: n_male_ = 5, n_female_ = 7) and 29 Tg (3m: n_male_ = 4, n_female_ = 4; 6m: n_male_ = 5, n_female_ = 6; 9m: n_male_ = 5, n_female_ = 5)

RSC containing slices were attached to a 0.1% poly-L-lysine coated coverslip and transferred to the recording chamber where they were perfused with carbogen-saturated aCSF (4 ml/min, 32-34°C). Differential interference contrast (DIC) and fluorescent signal imaging was performed using an Olympus BX51W1 microscope and SciCam Pro camera with a CoolLED pE-4000 LED light source. Borosilicate glass microelectrodes (OD 1.5mm, ID 0.86mm, 3-6 MΩ) were fabricated using a P-97 Flaming Brown micropipette puller and filled with caesium methanesulphonate (CsMeSO_4_) intracellular solution containing (in mM): 135 CsMeSO_4_, 8 KCl, 0.5 EGTA, 10 HEPES, 0.5 QX-314, 0.1 Spermine, 0.3 Na_2_-GTP, 2 Mg-ATP. Putative pyramidal cells (PC) were identified under DIC visualisation and assigned as being located in the superficial (L2-4) or deep (L5-6) layers of the dRSC or gRSC sub-regions of the RSC dependent on surrounding cytoarchitecture.

Data were collected with a Multiclamp 700B amplifier combined with a Digidata 1440A analogue-to-digital converter and a standalone computer equipped with pClamp software (Molecular Devices). Signals were digitised at 20 kHz and lowpass filtered at 8 kHz. Synaptic properties were recorded in voltage clamp mode, and cells were recorded from at a holding membrane potential (V_H_) of -70 mV and in standard aCSF unless otherwise stated. All optic stimulation was carried out at a 470 nm wavelength generated by a CoolLED pe-4000 light source at 100% power.

Cell response and EPSC magnitude were measured using a stimulation train protocol consisting of a 500 ms -10 mV square voltage injection followed by 5 ms optic stimulation pulses at 7 Hz frequency applied to the cell for 10-20 sweeps. PPR was measured using an increasing inter-pulse interval (IPI) protocol which consisted of: a 500 ms -10 mV square voltage injection followed by two 5 ms optic stimulation pulses at 10, 17, 51, 100, 170, 510 and 1000 ms intervals. Finally, AMPAR- and NMDAR-mediated currents were measured using a protocol consisting of a 500 ms -10 mV square voltage injection followed by a single 5 ms optic stimulation pulse. AMPAR-mediated currents were measured at V_H_ = -70 mV in standard aCSF and NMDAR-mediated currents were measured at V_H_ = +40 mV in aCSF containing antagonists for GABA_A_, GABA_B_ and non-NMDA ionotropic glutamate receptors (standard aCSF containing (in µM): 10 Gabazine, 1 CGP-55845, 10 DNQX).

Custom MATLAB scripts were used to analyse electrophysiological recordings unless stated otherwise. Electrophysiology traces were analysed using Igor Pro 7 (Wavemetrics. OR, USA) or MATLAB (Mathworks, CA, USA) software.

### Statistical Analysis

We used R^24^, implemented in RStudio^25^, to conduct all statistical analyses and generate all data graphs, using the lme4^26^, lmerTest^27^, tidyverse^28^, ggplot2^29^ and ggpubr packages.

## Results

We first confirmed that heterozygous hAPP-J20 mice showed increased deposition of Aβ with age. While no plaques were detectable at 3 months of age using Amylo-Glo^22^, Aβ expression was present at the other experimental time points, 6 and 9 months in the RSC (Supplementary Figure 1) and in both the hippocampus and the lateral EC (Supplementary Figure 2).

### Basal c-Fos reactivity is unchanged in hAPP-J20 mice

c-Fos is a widely-used marker of neuronal activation and it has been reported that its expression in RSC is reduced in the Tg2576 model of amyloidopathy before the onset of plaque deposition^16^. Furthermore, basal levels of c-Fos have been reported to be reduced in the dentate gyrus of hAPP-J20 mice at 4 to 6 months^30^ despite these mice displaying a hyperexcitability phenotype^31^. We carried out basal c-Fos staining in the RSC (Figure 1) as well as the CA1 region of the HPC and the EC (Figure 2) at 3, 6 and 9 months to determine whether these mice displayed signs of reduced neuronal activation.

**Figure 1:**
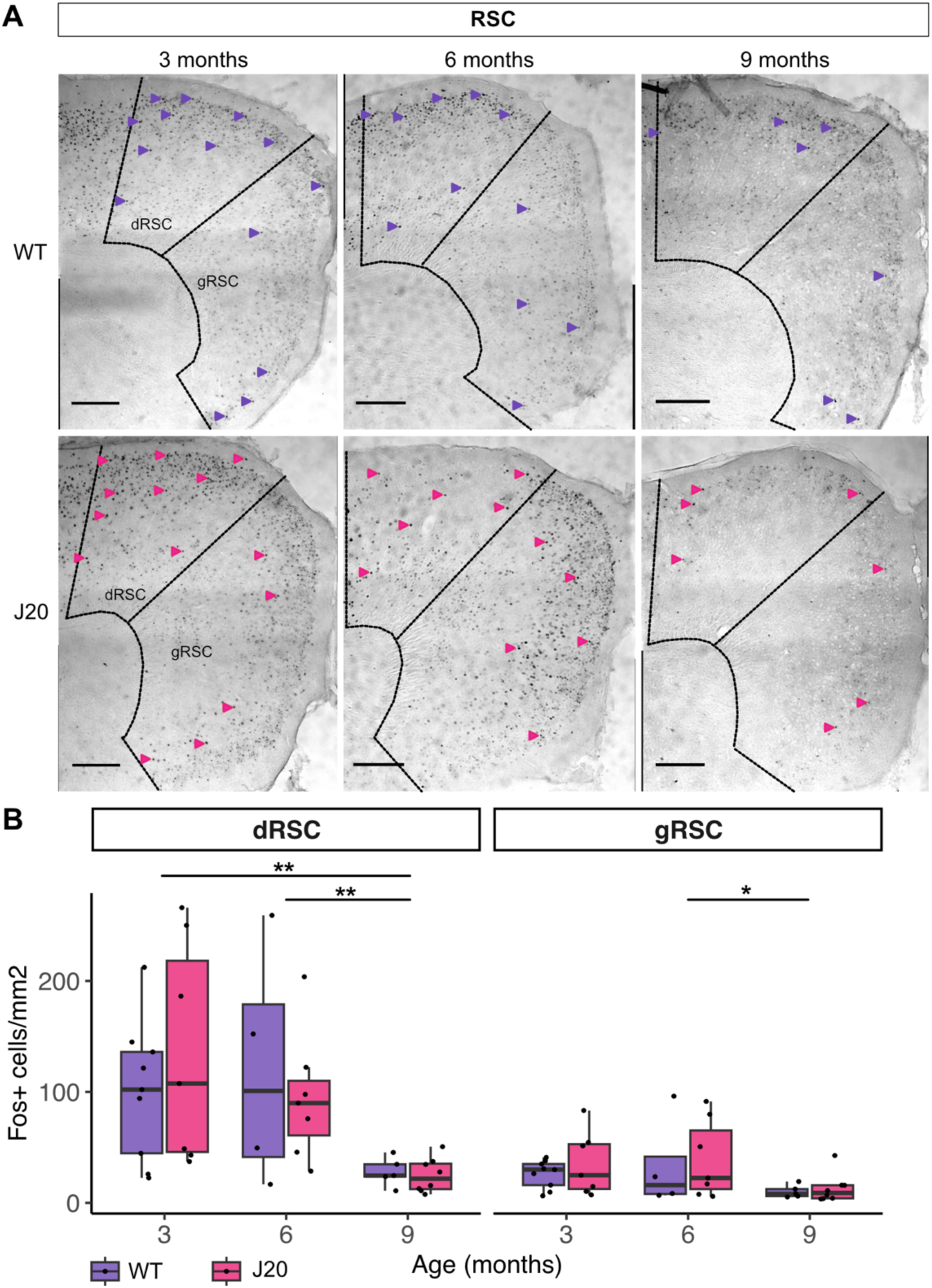
Basal c-Fos expression differs between RSC sub-regions and decreases with age but is not affected by hAPP-J20 genotype. **A,** representative images of basal c-Fos expression in the RSC in WT and Tg hAPP-J20 mice at 3m (WT, n=9; hAPP-J20, n=7), 6m (WT, n=4; hAPP-J20 n=7) and 9m (WT, n=5; hAPP-J20, n=8) age points. Images of the dRSC and gRSC were taken separately (10x magnification) and stitched together. Arrows indicate representative cells considered c-Fos^+^ but are not exhaustive of the count. Scale bars: 250 µm. **B,** a three-way mixed ANOVA indicated a significant main effect of sub-region (*F*(1,34) = 50.82, *p* < .001) and age (*F*(2,34) = 6.51, *p* < .01), as well as a significant interaction between the two variables (*F*(2, 34) = 7.86, *p* < .01). *Post hoc* tests comparing age points in the dRSC found significantly fewer c-Fos^+^ cells^mm2^ at 9m than 3m and 6m, which did not differ from each other (*p* = .65) (3m WT: 100.44 (62.26); 3m Tg: 134.15 (99.37); 6m WT: 119.39 (109.6); 6m Tg: 94.83 (57.47); 9m WT: 27.89 (12.97); 9m Tg: 24.69 (15.32) c-Fos^+^ cells^mm2^). In the gRSC, there were significantly fewer c-Fos^+^ cells^mm2^ at 9m than 6m, but not compared to 3m (*p* = .07). 3m and 6m also did not significantly differ in Fos expression (*p* = .43) (3m WT: 25.88 (12.44); 3m Tg: 35.17 (28.49); 6m WT: 33.73 (42.29); 6m Tg: 39.28 (35.08); 9m WT: 10.21 (5.73); 9m Tg: 12.97 (13.05) c-Fos^+^ cells^mm2^). There was no significant main effect of genotype (*F*(1, 34) = 0.08, *p* = .79), or interaction between genotype and age (*F*(2,34) = 0.45, *p* = .64) or sub-region (*F*(1,34) = 0.06, *p* = .81). Finally, there was no significant interaction between all three independent variables (*F*(2,34) = 0.99, *p* = .38). Descriptive stats display mean and SD. Boxplots display median (solid line), IQR and range. *\* p < .05, ** p < .01*

**Figure 2:**
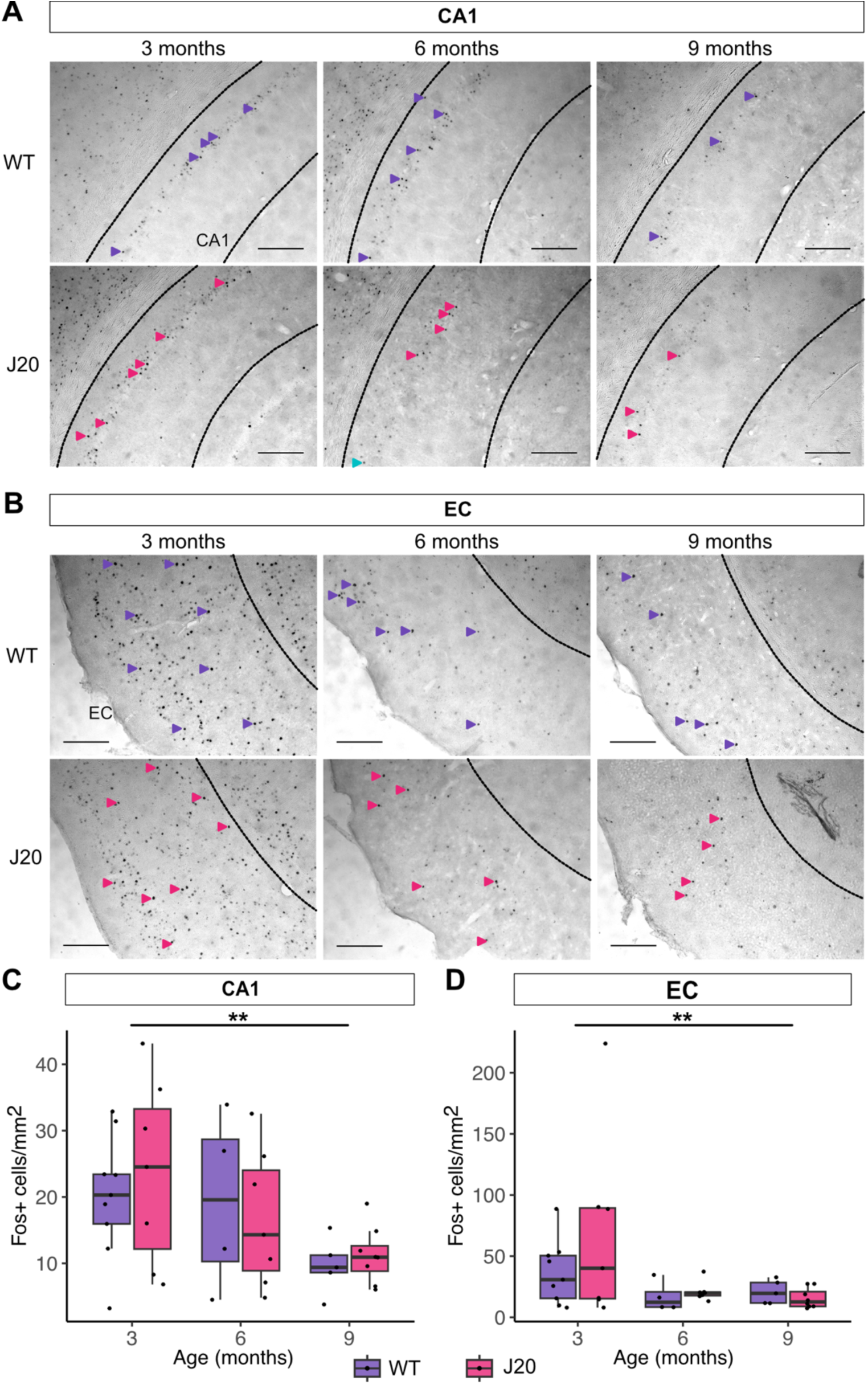
Basal c-Fos expression in CA1 and EC decreases with age but is not affected by hAPP-J20 genotype. **A-B,** Representative images of basal c-Fos expression in the CA1 (**A**) and EC (**B**) in WT and Tg hAPP-J20 mice at 3m, 6m and 9m age points. Images were taken at 10x magnification, and arrows indicate representative cells considered c-Fos^+^ but are not exhaustive of the count. Scale bars: 250 µm. **C,** in CA1 there was a main effect of age (*F*(2,34) = 4.75, *p* < .05; 2-way ANOVA), but no effect of genotype (*F*(1,34) = 0.002, *p* = .96) or interaction between age and genotype (*F*(2,34) = 0.30, *p* = .74). *Post hoc* tests found c-Fos expression was significantly lower at 9m than 3m, but there was no difference between 6m and 9m (*p* = .11) or 3m and 6m (*p* = .29) (**3m WT:** 20.18 (9.21); **3m Tg:** 23.62 (13.90); **6m WT:** 19.40 (13.43); **6m Tg:** 16.78 (10.35); **9m WT:** 9.67 (4.19); **9m Tg:** 11.22 (4.24) c-Fos^+^ cells^mm2^). **D,** in the EC there was a significant main effect of age (*F*(2,34) = 4.46, *p* < .05; 2-way ANOVA), but no main effect of genotype (*F*(1,34) = 0.33, *p* = .57) or interaction between age and genotype (*F*(2,34) = 0.11, *p* = .90). *Post hoc* tests indicated that c-Fos expression at 3m was significantly higher than 6m or 9m, while expression did not significantly differ between those age points (*p* = .89) (**3m WT:** 36.41 (26.00); **3m Tg:** 68.76 (76.55); **6m WT:** 16.85 (12.45); **6m Tg:** 20.98 (7.67); **9m WT:** 20.77 (9.63); **9m Tg:** 15.47 (8.17) c-Fos+ cells/mm^2^). Descriptive stats display mean and SD. Boxplots display median (solid line), IQR and range. *\*\* p < .01*.

We observed that basal c-Fos expression varied significantly been RSC subdivisions, with the dysgranular subdivision having a significantly higher level of expression than the granular subdivision (Figure 1B). We found that, while basal c-Fos expression was reduced at 9 months in both RSC subdivisions, there was no effect of genotype at any time point, with hAPP-J20 mice having comparable levels of basal c-Fos expression at 3, 6 and 9 months of age. No amyloid plaques were observed at 3 months but are apparent from 6 months of age in Tg mice, but not WT controls (Supplementary Figure 1). Furthermore, whilst c-Fos expression decreased with age, there was also a significant interaction between age and sub-region: the dRSC displayed a greater decrease in c-Fos+ cells as age increased. *Post hoc* analyses (Figure 1B) found no significant differences between WT and Tg mice at any age point in the dRSC (3m WT vs Tg: *p* = .42; 6m WT *vs* Tg: *p* = .63; 9m WT *vs* Tg: *p* = .71) or gRSC (3m WT *vs* Tg: *p* = .39; 6m WT *vs* Tg: *p* = .82; 9m WT *vs* Tg: *p* = .67). This suggests that while baseline neuronal activity is significantly decreased by 9 months (with a more pronounced reduction in dRSc with age), amyloidopathy does not alter basal c-Fos expression in this region at prodromal or clinical stages.

To examine whether this lack of effect of amyloidopathy on basal activity was restricted to the RSC, sections containing the CA1 (Figure 2A) and EC (Figure 2B) were also quantified and analysed. Both brain regions showed a similar pattern to the dRSC and gRSC in that c-Fos expression decreased with age, but was not affected by genotype (Figure 2C & D). *Post hoc* analyses in CA1 revealed no significant differences between wild-type and transgenic mice at each time point (3m WT *vs* Tg: *p* = .56; 6m WT *vs* Tg: *p* = .73; 9m WT *vs* Tg: *p* = .53), and the same was true in the EC (3m WT *vs* Tg: *p* = .25; 6m WT *vs* Tg: *p* = .51; 9m WT *vs* Tg: *p* = .31). Overall, we conclude that while amyloid plague deposition is occurring in these mice, and despite our previous observations that deficits in spatial memory occur in this line at 9 months of age^32^, there was no observable change in basal c-Fos expression in brain regions associated with spatial memory.

### No change in thalamic projections to RSC in hAPP-J20 mice

Deficits in synaptic transmission have been widely reported in hAPP-J20 mouse line^31,33–35^ as well as other amyloidopathy models. We recently reported that the ATN input to RSC is ubiquitous, with all excitatory neurons receiving input^36^. Given that the ATN have been targeted in attempts to reverse deficits in rodent models of dementia^32^ and other models of memory disorder^37^, we sought to test the hypothesis that ATN inputs to RSC would show age-dependent impairments in the hAPP-J20 mouse model. To do this, we carried out stereotaxic injections of AAV vectors into WT and hAPP-J20 mice to transduce expression of channelrhodospin into the ATN and carried out *ex vivo* patch-clamp electrophysiological recordings a 3, 6 and 9 months.

We targeted pyramidal neurons from deep and superficial layers of both dRSC and gRSC, recording from a total of 438 neurons, of which 305 passed quality checks (see methods), from 68 mice of both sexes. First, we checked for the presence of an EPSC in response to optogenetic stimulation of ATN axons. Like our previous findings in C57BL/6J mice^36^, all putative pyramidal cells responded to stimulation of this pathway for both genotypes and all age points. Therefore, probability of synaptic connectivity was not altered by age or genotype. Further analysis of the properties of the ATN-driven EPSC were then carried out (details in Table 1).

**Table 1.** Summary of all recorded and analysed neurons separated by age, genotype, RSC sub-region and cortical layer. All putative RSC pyramidal cells responded to stimulation of ATN afferent terminals. EPSC magnitude, onset time and 20-80% rise time were calculated from 305 neurons total. NMDA/AMPA ratio was calculated from 127 neurons of these cells, and paired pulse ratio (PPR) from 249 cells.

| Age | Genotype | Sub-region | Layer | Count |  |  |
| --- | --- | --- | --- | --- | --- | --- |
|  |  |  |  | EPSC measures | PPR | NMDA/AMPA |
| 3m (n=88) | WT (n=47) | dRSC | Superficial | 12 | 9 | 6 |
|  |  |  | Deep | 12 | 12 | 3 |
|  |  | gRSC | Superficial | 11 | 7 | 5 |
|  |  |  | Deep | 12 | 9 | 2 |
|  | Tg (n=41) | dRSC | Superficial | 10 | 9 | 5 |
|  |  |  | Deep | 8 | 7 | 3 |
|  |  | gRSC | Superficial | 11 | 10 | 5 |
|  |  |  | Deep | 12 | 7 | 2 |
| 6m (n=125) | WT (n=65) | dRSC | Superficial | 15 | 10 | 8 |
|  |  |  | Deep | 15 | 13 | 2 |
|  |  | gRSC | Superficial | 19 | 17 | 7 |
|  |  |  | Deep | 16 | 14 | 6 |
|  | Tg (n=60) | dRSC | Superficial | 20 | 16 | 10 |
|  |  |  | Deep | 13 | 11 | 7 |
|  |  | gRSC | Superficial | 13 | 11 | 7 |
|  |  |  | Deep | 14 | 12 | 8 |
| 9m (n=92) | WT (n=56) | dRSC | Superficial | 13 | 12 | 8 |
|  |  |  | Deep | 16 | 13 | 6 |
|  |  | gRSC | Superficial | 12 | 11 | 6 |
|  |  |  | Deep | 15 | 15 | 6 |
|  | Tg (n=36) | dRSC | Superficial | 13 | 8 | 7 |
|  |  |  | Deep | 5 | 3 | 1 |
|  |  | gRSC | Superficial | 12 | 9 | 5 |
|  |  |  | Deep | 6 | 4 | 2 |
| Total |  |  |  | 305 | 249 | 127 |

To examine the magnitude of the AMPA receptor-mediated component of the ATN-EPSC onto RSC pyramidal cells, a null model (*random effect*: mouse) and three mixed effect models were created. The goodness-of-fit (represented by AIC) was sequentially evaluated using χ^2^ likelihood ratio comparisons. Model 1 (*random effect*: mouse; *fixed effects:* sub-region and layer) significantly improved upon the null model (*χ_2_*(2) = 73.0, *p* < .001; AIC_null_ = 4595.0, AIC_M1_ = 4526.0), and both sub-region (*F*(1,280.1) = 62.2, *p* < .001; ANOVA) and layer (*F*(1,286.1) = 286.1, *p* < .001; ANOVA) significantly affected EPSC magnitude. Fixed effect estimates (Table 2) indicate that cells in the gRSC and deep layers had a significantly lower EPSC magnitude compared to those in the dRSC and superficial layers, respectively. Inspection of the data shows that cells in the superficial layers of the dRSC had the largest EPSC magnitude, followed by dRSC deep layer cells (Figure 3) in line with our previous findings^36^ . Next, model 2 added age and genotype as fixed effects (*random effect:* mouse; *fixed effects*: sub-region, layer, age and genotype); however this model did not improve upon model 1 (*χ_2_*(3) = 3.9, *p* = .27; AIC_M2_ = 4528.1). Furthermore, the addition of sex as a fixed effect in model 3 (*random effect:* mouse; *fixed effects*: sub-region, layer, age, genotype and sex) did not improve upon model 2 (*χ_2_*(1) = 2.8, *p* = 0.09; AIC_M3_ = 4527.7), and fixed effect estimates for models 2 and 3 did not show any significant differences in EPSC magnitude for these added factors (Table 2). Overall, model 1 was the best model of EPSC magnitude variability, suggesting age, genotype and sex have no effect on this measure.

**Figure 3:**
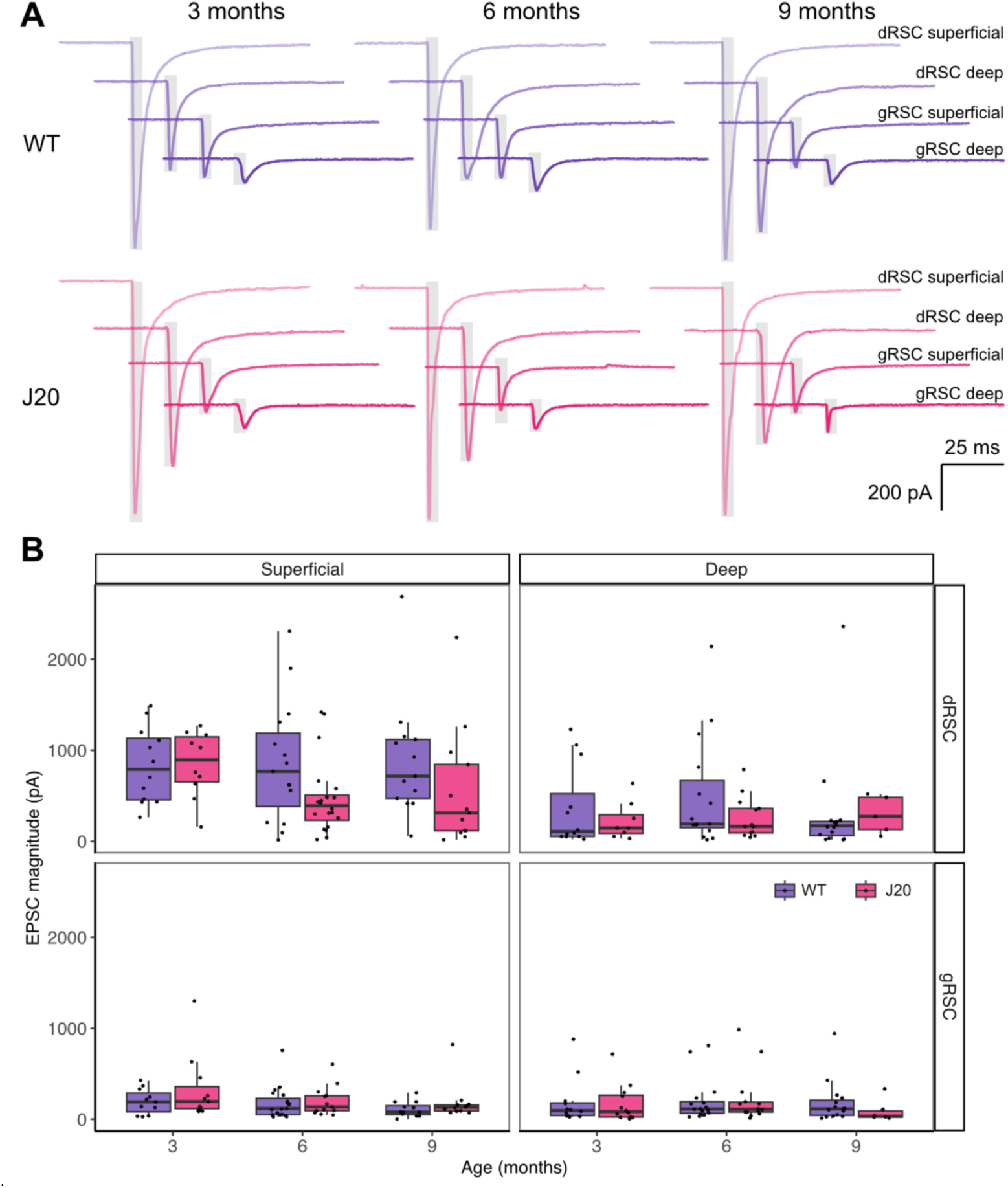
EPSC magnitude differs between RSC sub-region and cortical layer, but is not affected by age or J20 genotype. **A,** Representative voltage-clamp (V_H_ = -70 mV) traces showing the first EPSC generated from a 7 Hz stimulation protocol. Grey boxes indicate the 5 ms optical stimulation period (470 nm wavelength). **B,** EPSC magnitude differed between RSC sub-region and cortical layer: magnitude appeared largest in the superficial layers of the dRSC, followed by the deep layers dRSC whilst gRSC superficial and deep layer EPSC magnitude appeared the smallest. Neither age nor genotype significantly explained variance in EPSC magnitude, and no clear difference in these factors are observed graphically. Boxplots display median (solid line), IQR and range.

**Table 2:**
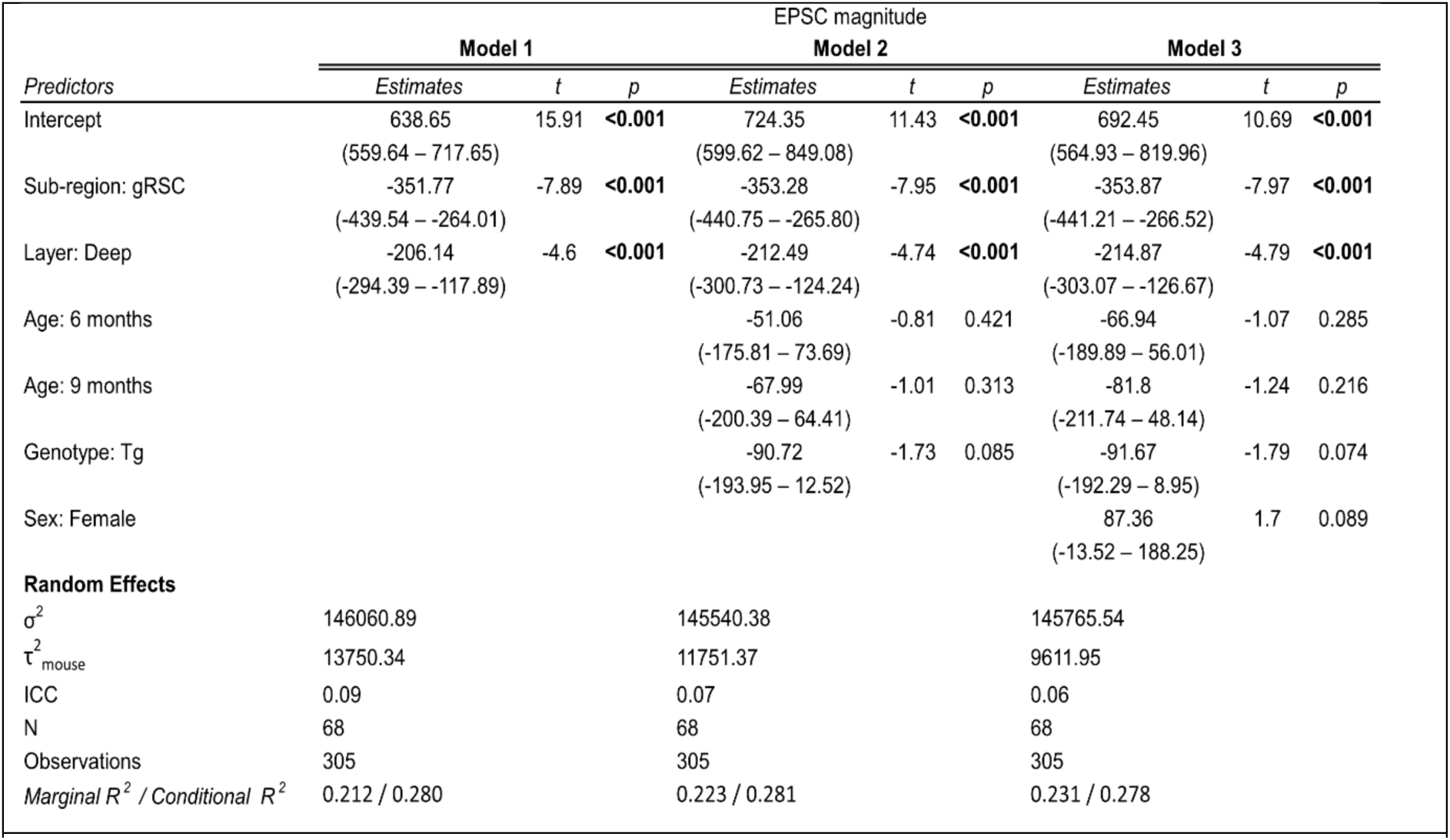
Fixed and random effect results for each mixed model analysing EPSC magnitude. For fixed effects (predictors), the Table displays effect size, confidence intervals, *t* statistic and significance value. For random effects, the Table displays residual variance (σ^2^), mouse variance (τ^2^) and the intra-class correlation coefficient (ICC). Marginal R^2^ refers to variance explained by fixed effects only, while conditional R^2^ refers to variance explained by combined fixed and random effects. There was also a random effect of mouse present in model 1, contributing 6.8% of variance to the total explained by the model, although the ICC score indicates a weak resemblance between mice

We next analysed the time of onset of EPSC after optogenetic stimulation and the 20 to 80% rise time of the EPSC to infer whether amyloidopathy perturbed presynaptic release synchrony (Supplementary Figure 3). Superficial pyramidal cells displayed a slightly faster EPSC onset than deeper pyramidal cells (Supplementary Figure 3A & B), which would be expected due to ATN axons entering RSC in superficial layers^36^. While modelling revealed that RSC subdivision, laminar location and sex had significant effects on the EPSC onset time, there was no effect of age or genotype with WT and hAPP-J20 mice displaying no differences (Supplementary Table 1). We found no effect of RSC subdivision or laminar location on 20-80% rise time, and subsequent models including age, genotype and sex did not improve upon the null model (Supplementary Figure 3C and Supplementary Table 2).

Our final analysis of the AMPA-mediated component of the ATN-EPSC onto RSC pyramidal cells was to examine the paired pulse ratio (PPR) to determine whether hAPP overexpression led to changes in short-term synaptic plasticity (Supplementary Figure 4) at different inter-stimulus intervals. As we reported previously^36^, PPR varied significantly between dSRC and gRSC, as well as between deep and superficial layers, but we observed no effect of age or genotype (Supplementary Figure 4 and Supplementary Table 4). Overall, we concluded that neither age up to 9 months nor the presence or absence of amyloidopathy had any significant effect on thalamo-retrosplenial signalling from the ATN.

### No change in NMDA / AMPA ratio in hAPP-J20 mice at any time point

Finally, we sought to determine whether the NMDA/AMPA ratio was altered in hAPP-J20 mice, as this is a sensitive measure of postsynaptic ionotropic glutamate receptor function and potential for plasticity at synapses. Example electrophysiological traces showing AMPA and NMDA-receptor mediated components of the ATN-EPSC are shown in Figure 4 for each age, genotype and RSC subdivision. Again, we used a series of linear mixed effects models to determine the influences of these factors on the NMDA/AMPA ratio (Table 3).

**Figure 4:**
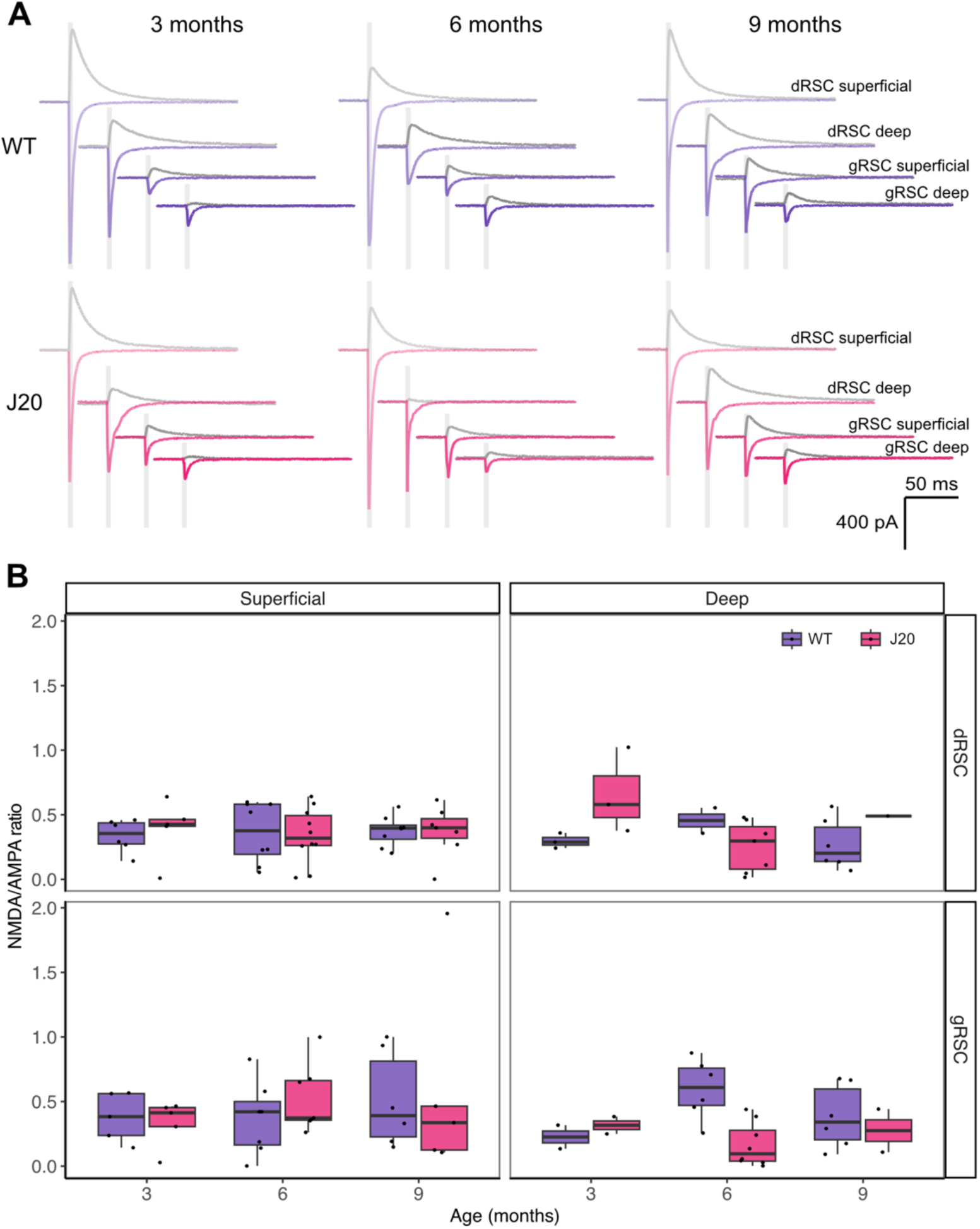
NMDA/AMPA ratio does not differ between sub-region cortical layer, age or hAPP-J20 genotype. **A,** representative voltage-clamp traces showing AMPAR-mediated (blue/red trace) and NMDAR-mediated EPSCs (grey trace) generated by optogenetic stimulation (grey box indicates 5 ms optical stimulation period). AMPAR responses were recorded at V_H_ = -70 mV (standard aCSF) and NMDAR responses were recorded at V_H_ = +40 mV (standard aCSF containing 10 µM DNQX, 1 µM CGP55845 and 10 µM gabazine). **B**, NMDA/AMPA ratio did not differ between sub-region, layer, age or genotype. Boxplots display median (solid line), IQR and range.

**Table 3:**
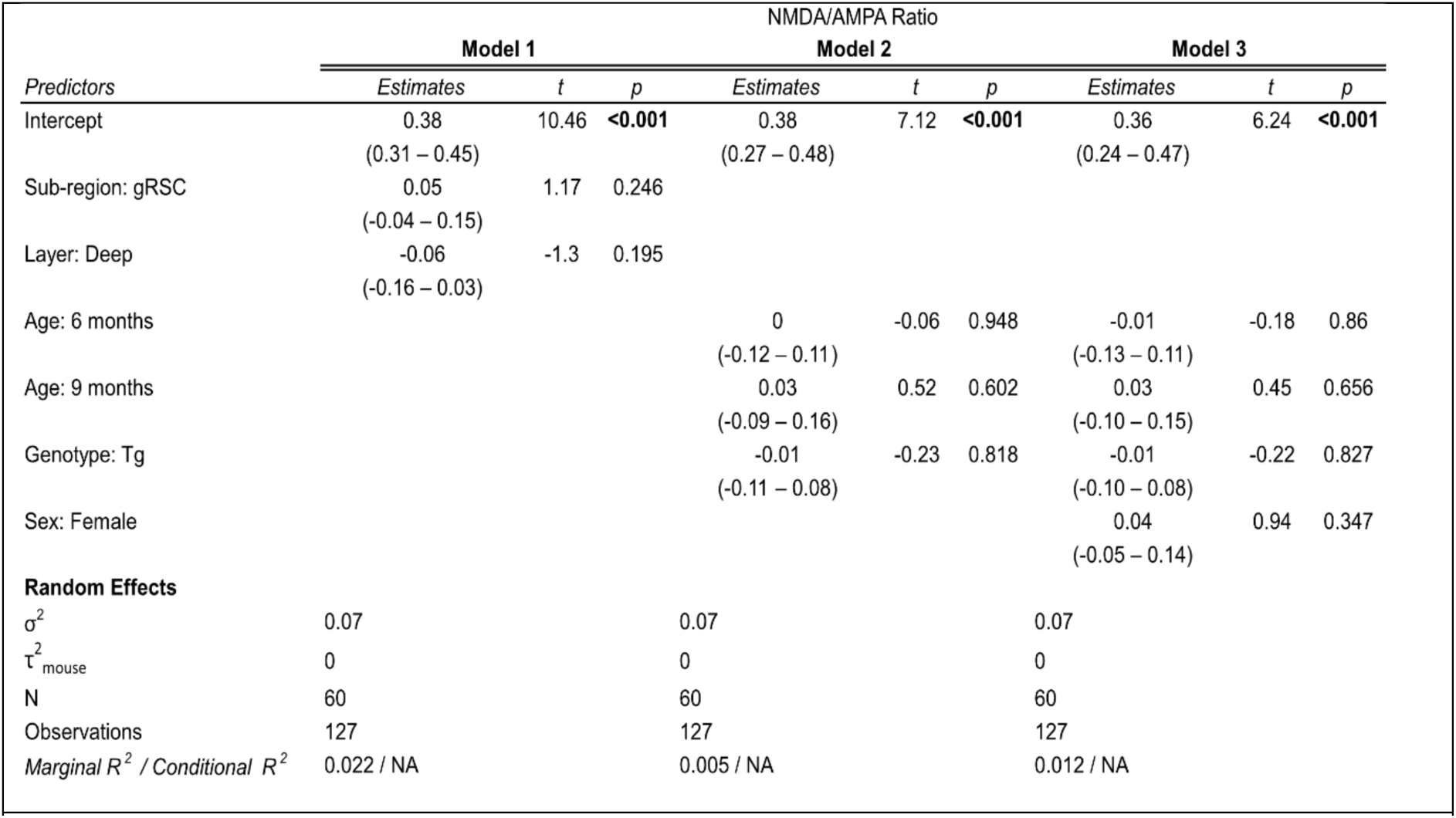
Fixed and random effect results for each mixed model analysing NMDA/AMPA ratio. For fixed effects (predictors), the Table displays effect size, confidence intervals, *t* statistic and significance value. For random effects, the Table displays residual variance (σ^2^), mouse variance (τ^2^) and the ICC value. Marginal R^2^ refers to variance explained by fixed effects only, while conditional R^2^ refers to variance explained by combined fixed and random effects.

Analysis of NMDA/AMPA ratios found that model 1 (*fixed effects:* sub-region and layer) did not significantly improve upon the null model (*χ_2_*(2) = 2.8, *p* = .25; AIC_null_ = 28.9, AIC_M1_ = 30.2), and fixed effect estimates indicated no significant differences in NMDA/AMPA ratio between sub-regions and layers (Table 4.5). Model 2 (*fixed effects:* age and genotype) omitted area and layer as factors due to the lack of significance in model 1, but again did improve upon the null model (*χ_2_*(3) = 0.6, *p* = .89; AIC_M2_ = 34.3). Moreover, the addition of sex as a factor to model 3 did not significantly improve on model 2 (*χ_2_*(1) = 0.9, *p* = .35; AIC_M3_ = 35.4). Fixed effect estimates for age, genotype and sex in models 2 and 3 found no significant differences within these factors (Table 3). Finally, the null model found a negligible random effect of mouse on variance, which was also true in models 1-3 (Table 3 random effects). Therefore NMDA/AMPA ratio was not significantly affected by sub-region, layer, age or genotype (Figure 4) and did not differ significantly between mice.

## Discussion

In this study, we set out to test the hypotheses that the RSC in hAPP-J20 mice would display evidence of the hypoactivity that is present in prodromal AD in humans^8,9^, and that specific deficits in projections from the ATN to the RSC would be present in this model. However, we found no evidence to support either of these hypotheses and can conclude that signalling through ATN, at least to the RSC, remains intact in the hAPP-J20 model, in contrast to deficits seen in areas such as hippocampus at similar time points in this model^31,33–35^. This surprising lack of change in this circuit helps to explain why we observed no rescue of spatial memory when increasing the excitability of ATN using chemogenetics in the hAPP-J20 mouse model, but could rescue novel object memory when targeting thalamic nucleus reuniens^32^.

### Basal c-Fos reactivity in the RSC

We found that the basal c-Fos reactivity was higher in dysgranular RSC compared with the granular subdivision. gRSC is associated with allocentric inputs arriving from dorsal subiculum^38^, although we found that subicular inputs could similarly excite neurons in both dRSC and gRSC^36^, while visual information associated with egocentric circuits preferentially arrives in dRSC^38^. Given that the mice used for basal c-Fos were housed in their home cage before perfusion, being in a familiar environment without the need to navigate may account for the lower c-Fos reactivity in gRSC, when dRSC would likely still be receiving input from the visual system irrespective of the behaviour of the mice.

Our observation that c-Fos expression in the RSC, CA1 and EC decreased with age is consistent with prior literature that describes decreased hippocampal and cortical basal levels of Fos protein and mRNA in aged rats^39,40^. The mechanism behind this decrease is not clear, however, it may be linked to ageing-induced impairments in upstream signalling pathways such as MAPK^41^, which is the primary pathway through which c-Fos expression is mediated^42^. While the number of Fos^+^ cells decreased with age, there were no differences in basal c-Fos expression between WT and hAPP-J20 mice at 3, 6 or 9 months of age. This was an unexpected finding, as this marker of neuronal activity has been shown to be reduced in the RSC in another model of Aβ pathology^16^. However, this reduction in c-Fos induction was observed following exposure to a novel environment. A different study using the APPswe/PS1ΔE9 model found reduced *Fos* mRNA expression in the CA1 region only after exploration of a novel environment but not in basal c-Fos levels^43^. It may be that not using a behavioural spatial memory, navigation or exploration in our task failed to engage the RSC sufficiently to reveal any potential differences in RSC activity between wildtype of hAPP-J20 mice. Alternatively, there may just be no deficit present in this model at the time points that we measured.

### Synaptic properties of ATN projections to the RSC

We have previously found that both midline and anterior thalamic projections to cortical regions in the extended memory circuit elicit strong EPSCs, with both the nucleus reuniens^44^ and the ATN^36^ driving currents in the hundreds to thousands of pA range. Indeed, we previously found that the ATN formed monosynaptic connections with every pyramidal cell that we recorded from in RSC^36^, and in the present study found every neuron out of the 305 that we recorded from received direct thalamic input. This study confirmed out previous findings^36^ that optogenetic stimulation of ATN axons drove larger EPSCs in dRSC than gRSC, and larger responses in superficial pyramidal cells. However, we found that this projection did not change with age and appeared unaffected by amyloidopathy in the hAPP-J20 mouse model. It should be noted that this study targeted glutamatergic neurons in RSC only, and it is possible that thalamic input to GABAergic neurons in RSC may be perturbed in this model, given the disruption to inhibitory neuron signalling observed by others in the hAPP-J20 line.^35^

As discussed earlier, the RSC is particularly sensitive to deafferentation^17,45^, and the ATN provide substantial glutamatergic input to this region. While the anterior thalamus does show amyloid pathology^1,2^, we did not observe any amyloid plaque deposition in the ATN in hAPP-J20 mice at any time point tested. We hypothesised that Aβ oligomers present in the RSC would have synaptotoxic effects sufficient to disrupt this pathway, as is seen in other models and brain regions^46–48^. However, our results indicate that post-synaptic Aβ pathology is not enough to impair synaptic responses and, therefore, suggests that disruption of the ATN to RSC pathway may not mediate cognitive deficits in pre-clinical and prodromal AD. This is supported by our recent findings that chemogenetic activation of the ATN does not improve behavioural deficits in J20 mice at 6m or 9m of age^32^.

RSC afferent disruption may still be involved in cognitive impairment in AD, as pathways other than those with the thalamus could be affected in the early stages of the disease. One possible candidate is the dorsal subiculum input, which also exerts a strong excitatory influence on both subdivisions of the RSC^36^. The dorsal subiculum contains a range of spatially-tuned cells and sends information on speed, trajectory and place to the RSC^49^. Not only does the dorsal subiculum develop plaques at similar age-points to the RSC in hAPP-J20 mice^50^, but lesioning this region also reduces the spread of Aβ pathology into the RSC^51^. The RSC receives inputs from many brain regions, but it also sends efferents to widely distributed brain regions. Therefore, as the RSC is one of the first regions to show Aβ pathology in humans, perhaps its connectivity dysfunction lies in its efferent outputs. It is also possible that other mouse models of AD could show disrupted connectivity between the ATN and RSC, and that our chosen model was an unfortunate outlier for studying the cellular connectivity between these brain regions in dementia.

## Acknowledgements

This work was supported by an Alzheimer’s Research UK Interdisciplinary research grant ARUK-IRG2017B-4 (MTC), and salary support for LA came from Biotechnology and Biological Sciences Research Council grant BB/P001475/1 (MTC). GMS was a GW4 BioMed doctoral training program student funded by the Medical Research Council (MR/N0137941/1). For the purpose of open access, the authors have applied a Creative Commons Attribution (CC-BY) licence to any Author Accepted Manuscript version arising from this submission.

## Author Contributions

Conceptualization, GM and MTC.; Methodology, GM, AR and MTC; Investigation, GM, LA, and SK, MTC; Funding Acquisition, MTC, AR, JPA; Resources, MTC and AR; Supervision, MTC, AR, JPA, and JW. All authors contributed to the final version of the manuscript.

## Data Availability

All data supporting the results are available within the manuscript. Numerical data will be deposited in our institutional repository upon acceptance and will be freely available.

## References

1. Braak H, Braak E. Neuropathological stageing of Alzheimer-related changes. Acta Neuropathol. 1991;82(4):239–259. doi:10.1007/bf00308809

2. Braak H, Braak E. Alzheimer’s disease affects limbic nuclei of the thalamus. Acta Neuropathol. 1991;81(3):261–268. doi:10.1007/bf00305867

3. Aggleton JP, Pralus A, Nelson AJD, Hornberger M. Thalamic pathology and memory loss in early Alzheimer’s disease: moving the focus from the medial temporal lobe to Papez circuit. Brain. 2016;139(7):1877–1890. doi:10.1093/brain/aww083

4. Barnes J, Dickerson BC, Frost C, Jiskoot LC, Wolk D, Flier WM van der. Alzheimer’s disease first symptoms are age dependent: Evidence from the NACC dataset. Alzheimer’s Dement. 2015;11(11):1349–1357. doi:10.1016/j.jalz.2014.12.007

5. Porsteinsson AP, Isaacson RS, Knox S, Sabbagh MN, Rubino I. Diagnosis of Early Alzheimer’s Disease: Clinical Practice in 2021. J Prev Alzheimer’s Dis. 2021;8(3):371–386. doi:10.14283/jpad.2021.23

6. Aggleton JP. Looking beyond the hippocampus: old and new neurological targets for understanding memory disorders. Proc Royal Soc B Biological Sci. 2014;281(1786):20140565. doi:10.1098/rspb.2014.0565

7. Vann SD, Aggleton JP, Maguire EA. What does the retrosplenial cortex do? Nat Rev Neurosci. 2009;10(11):792–802. doi:10.1038/nrn2733

8. Nestor PJ, Fryer TD, Ikeda M, Hodges JR. Retrosplenial cortex (BA 29/30) hypometabolism in mild cognitive impairment (prodromal Alzheimer’s disease). Eur J Neurosci. 2003;18(9):2663–2667. doi:10.1046/j.1460-9568.2003.02999.x

9. Dillen KNH, Jacobs HIL, Kukolja J, et al. Aberrant functional connectivity differentiates retrosplenial cortex from posterior cingulate cortex in prodromal Alzheimer’s disease. Neurobiol Aging. 2016;44:114–126. doi:10.1016/j.neurobiolaging.2016.04.010

10. Palmqvist S, Schöll M, Strandberg O, et al. Earliest accumulation of β-amyloid occurs within the default-mode network and concurrently affects brain connectivity. Nat Commun. 2017;8(1):1214. doi:10.1038/s41467-017-01150-x

11. Pengas G, Hodges JR, Watson P, Nestor PJ. Focal posterior cingulate atrophy in incipient Alzheimer’s disease. Neurobiol Aging. 2010;31(1):25–33. doi:10.1016/j.neurobiolaging.2008.03.014

12. Scheff SW, Price DA, Ansari MA, et al. Synaptic Change in the Posterior Cingulate Gyrus in the Progression of Alzheimer’s Disease. J Alzheimer’s Dis. 2014;43(3):1073–1090. doi:10.3233/jad-141518

13. Villain N, Desgranges B, Viader F, et al. Relationships between Hippocampal Atrophy, White Matter Disruption, and Gray Matter Hypometabolism in Alzheimer’s Disease. J Neurosci. 2008;28(24):6174–6181. doi:10.1523/jneurosci.1392-08.2008

14. Terstege DJ, Ren Y, Ahn BY, et al. Impaired parvalbumin interneurons in the retrosplenial cortex as the cause of sex-dependent vulnerability in Alzheimer’s disease. Sci Adv. 2025;11(18):eadt8976. doi:10.1126/sciadv.adt8976

15. Kim DH, Kim HA, Han YS, Jeon WK, Han JS. Recognition memory impairments and amyloid-beta deposition of the retrosplenial cortex at the early stage of 5XFAD mice. Physiol Behav. 2020;222:112891. doi:10.1016/j.physbeh.2020.112891

16. Poirier GL, Amin E, Good MA, Aggleton JP. Early-onset dysfunction of retrosplenial cortex precedes overt amyloid plaque formation in Tg2576 mice. Neuroscience. 2011;174:71–83. doi:10.1016/j.neuroscience.2010.11.025

17. Albasser MM, Poirier GL, Warburton EC, Aggleton JP. Hippocampal lesions halve immediate–early gene protein counts in retrosplenial cortex: distal dysfunctions in a spatial memory system. Eur J Neurosci. 2007;26(5):1254–1266. doi:10.1111/j.1460-9568.2007.05753.x

18. Mucke L, Masliah E, Yu GQ, et al. High-Level Neuronal Expression of Aβ1–42 in Wild-Type Human Amyloid Protein Precursor Transgenic Mice: Synaptotoxicity without Plaque Formation. J Neurosci. 2000;20(11):4050–4058. doi:10.1523/jneurosci.20-11-04050.2000

19. Paxinos G, Franklin KBJ. Paxinos and Franklin’s the Mouse Brain in Stereotaxic Coordinates.; 2001. Accessed January 19, 2022. https://www.elsevier.com/books/paxinos-and-franklins-the-mouse-brain-in-stereotaxic-coordinates/paxinos/978-0-12-816157-9

20. Schindelin J, Arganda-Carreras I, Frise E, et al. Fiji: an open-source platform for biological-image analysis. Nat Methods. 2012;9(7):676–682. doi:10.1038/nmeth.2019

21. Bolte S, Cordelières FP. A guided tour into subcellular colocalization analysis in light microscopy. J Microsc. 2006;224(3):213–232. doi:10.1111/j.1365-2818.2006.01706.x

22. Schmued L, Raymick J, Tolleson W, Sarkar S, Zhang YH, Bell-Cohn A. Introducing Amylo-Glo, a novel fluorescent amyloid specific histochemical tracer especially suited for multiple labeling and large scale quantification studies. J Neurosci Methods. 2012;209(1):120–126. doi:10.1016/j.jneumeth.2012.05.019

23. Ting JT, Daigle TL, Chen Q, Feng G. Acute brain slice methods for adult and aging animals: application of targeted patch clamp analysis and optogenetics. Methods Mol Biol. 2014;1183:221–242. doi:10.1007/978-1-4939-1096-0_14

24. Team RC. R: A Language and Environment for Statistical Computing. Published online 2021. https://www.R-project.org/

25. Team RStudio. RStudio: Integrated Development for R. RStudio.; 2020. http://www.rstudio.com/

26. Bates D, Mächler M, Bolker B, Walker S. Fitting Linear Mixed-Effects Models Using lme4. J Stat Softw. 2015;67(1). doi:10.18637/jss.v067.i01

27. Kuznetsova A, Brockhoff PB, Christensen RHB. lmerTest Package: Tests in Linear Mixed Effects Models. J Stat Softw. 2017;82(13). doi:10.18637/jss.v082.i13

28. Wickham H, Averick M, Bryan J, et al. Welcome to the tidyverse. Journal of Open Source Software. 2019;4(43):1686. doi:10.21105/joss.01686

29. Wickham H. ggplot2, Elegant Graphics for Data Analysis. Published online 2016:11–31. doi:10.1007/978-3-319-24277-4_2

30. Corbett BF, You JC, Zhang X, et al. ΔFosB Regulates Gene Expression and Cognitive Dysfunction in a Mouse Model of Alzheimer’s Disease. Cell Rep. 2017;20(2):344–355. doi:10.1016/j.celrep.2017.06.040

31. Palop JJ, Chin J, Roberson ED, et al. Aberrant Excitatory Neuronal Activity and Compensatory Remodeling of Inhibitory Hippocampal Circuits in Mouse Models of Alzheimer’s Disease. Neuron. 2007;55(5):697–711. doi:10.1016/j.neuron.2007.07.025

32. Kohli S, Andrianova L, Margetts-Smith G, Brady E, Craig MT. Chemogenetic activation of midline thalamic nuclei fails to ameliorate memory deficits in two mouse models of Alzheimer’s disease. bioRxiv. Published online July 3, 2021. doi:10.1101/2021.06.30.450500

33. Sun B, Zhou Y, Halabisky B, et al. Cystatin C-Cathepsin B Axis Regulates Amyloid Beta Levels and Associated Neuronal Deficits in an Animal Model of Alzheimer’s Disease. Neuron. 2008;60(2):247–257. doi:10.1016/j.neuron.2008.10.001

34. Harris JA, Devidze N, Halabisky B, et al. Many Neuronal and Behavioral Impairments in Transgenic Mouse Models of Alzheimer’s Disease Are Independent of Caspase Cleavage of the Amyloid Precursor Protein. J Neurosci. 2010;30(1):372–381. doi:10.1523/jneurosci.5341-09.2010

35. Verret L, Mann EO, Hang GB, et al. Inhibitory Interneuron Deficit Links Altered Network Activity and Cognitive Dysfunction in Alzheimer Model. Cell. 2012;149(3):708–721. doi:10.1016/j.cell.2012.02.046

36. Margetts-Smith G, Andrianova L, Kohli S, et al. Dissection of retrosplenial cortex inputs: ubiquitous drive from anterior thalamus. bioRxiv. Published online February 14, 2025:2025.02.06.636939. doi:10.1101/2025.02.06.636939

37. Barnett SC, Parr-Brownlie LC, Perry BAL, et al. Anterior thalamic nuclei neurons sustain memory. Curr Res Neurobiology. 2021;2:100022. doi:10.1016/j.crneur.2021.100022

38. Aggleton JP, Yanakieva S, Sengpiel F, Nelson AJ. The separate and combined properties of the granular (area 29) and dysgranular (area 30) retrosplenial cortex. Neurobiol Learn Mem. 2021;185:107516. doi:10.1016/j.nlm.2021.107516

39. Kitraki E, Bozas E, Philippdis H, Stylianopoulou F. Aging-related changes in IGF-II and c-fos gene expression in the rat brain. Int J Dev Neurosci. 1993;11(1):1–9. doi:10.1016/0736-5748(93)90029-d

40. Lee YI, Park KH, Baik SH, Cha CI. Attenuation of c-Fos basal expression in the cerebral cortex of aged rat. NeuroReport. 1998;9(12):2733–2736. doi:10.1097/00001756-199808240-00009

41. Zhen X, Uryu K, Cai G, Johnson GP, Friedman E. Age-Associated Impairment in Brain MAPK Signal Pathways and the Effect of Caloric Restriction in Fischer 344 Rats. J Gerontol Ser A: Biomed Sci Méd Sci. 1999;54(12):B539–B548. doi:10.1093/gerona/54.12.b539

42. Chung L. A Brief Introduction to the Transduction of Neural Activity into Fos Signal. Dev Reprod. 2015;19(2):61–67. doi:10.12717/dr.2015.19.2.061

43. Christensen DZ, Thomsen MS, Mikkelsen JD. Reduced basal and novelty-induced levels of activity-regulated cytoskeleton associated protein (Arc) and c-Fos mRNA in the cerebral cortex and hippocampus of APPswe/PS1ΔE9 transgenic mice. Neurochem Int. 2013;63(1):54–60. doi:10.1016/j.neuint.2013.04.002

44. Andrianova L, Banks PJ, Booth CA, et al. Hippocampal pyramidal cells of the CA1 region are not a major target of the thalamic nucleus reuniens. PLOS Biol. 2025;23(10):e3003419. doi:10.1371/journal.pbio.3003419

45. Dumont JR, Amin E, Poirier GL, Albasser MM, Aggleton JP. Anterior thalamic nuclei lesions in rats disrupt markers of neural plasticity in distal limbic brain regions. Neuroscience. 2012;224:81–101. doi:10.1016/j.neuroscience.2012.08.027

46. Hsia AY, Masliah E, McConlogue L, et al. Plaque-independent disruption of neural circuits in Alzheimer’s disease mouse models. Proc Natl Acad Sci. 1999;96(6):3228–3233. doi:10.1073/pnas.96.6.3228

47. Shipton OA, Leitz JR, Dworzak J, et al. Tau Protein Is Required for Amyloid β-Induced Impairment of Hippocampal Long-Term Potentiation. J Neurosci. 2011;31(5):1688–1692. doi:10.1523/jneurosci.2610-10.2011

48. Shipton OA, Tang CS, Paulsen O, Vargas-Caballero M. Differential vulnerability of hippocampal CA3-CA1 synapses to Aβ. Acta Neuropathol Commun. 2022;10(1):45. doi:10.1186/s40478-022-01350-7

49. Kitanishi T, Umaba R, Mizuseki K. Robust information routing by dorsal subiculum neurons. Sci Adv. 2021;7(11):eabf1913. doi:10.1126/sciadv.abf1913

50. Whitesell JD, Buckley AR, Knox JE, et al. Whole brain imaging reveals distinct spatial patterns of amyloid beta deposition in three mouse models of Alzheimer’s disease. J Comp Neurol. 2019;527(13):2122–2145. doi:10.1002/cne.24555

51. George S, Rönnbäck A, Gouras GK, et al. Lesion of the subiculum reduces the spread of amyloid beta pathology to interconnected brain regions in a mouse model of Alzheimer’s disease. Acta Neuropathol Commun. 2014;2(1):17. doi:10.1186/2051-5960-2-17

