## Supplementary material for "Anterior thalamic nucleus inputs to the retrosplenial cortex are unperturbed in the hAPP-J20 mouse model of Alzheimer’s disease"

**Supplementary figures**

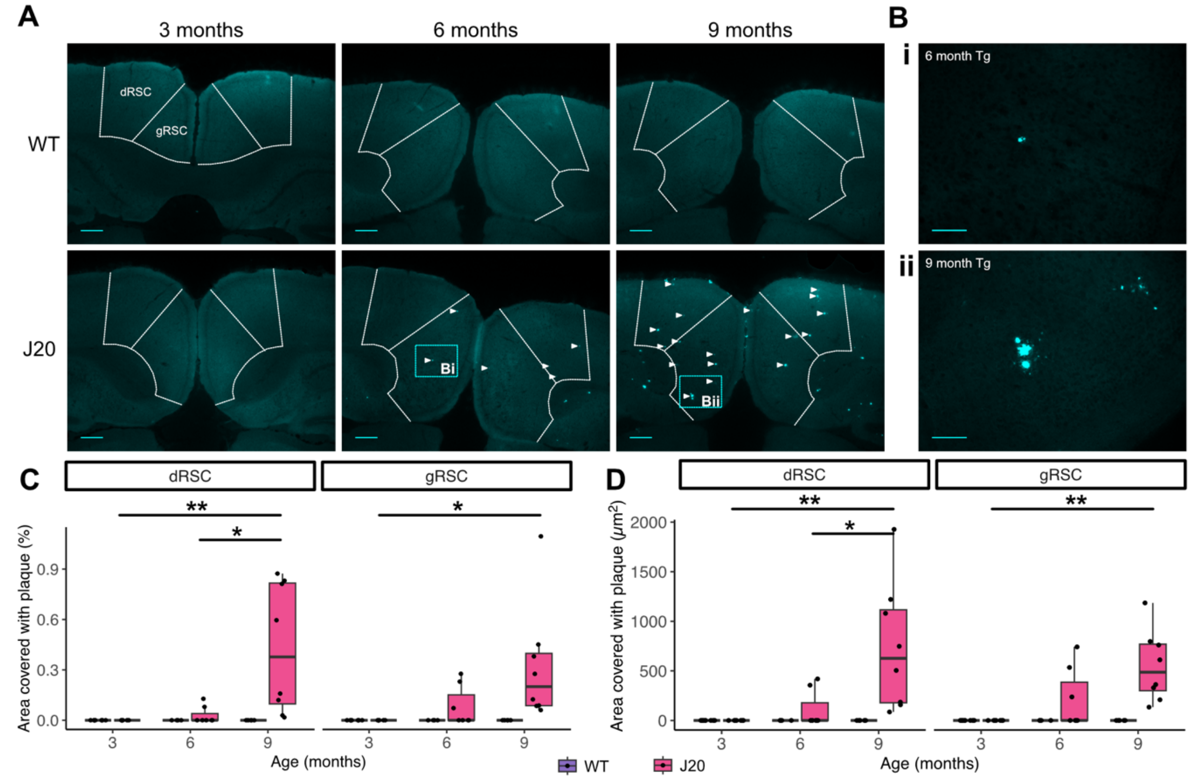


**Supplementary figure 1: amyloid plaque deposition in retrosplenial cortex increases with age in J20 mice. A,** representative images of Aβ plaque deposition in the RSC in WT and J20 mice at 3m, 6m and 9m. Arrows indicate plaques at 6m and 9m in J20 mice. Scale bars: 250 μm. **B,** higher magnification images displaying representative plaques from the 6m (**Bi**) and 9m (**Bii**) J20 mice sections respectively. Scale bars: 100 μm. **C,** analysis of dRSC and gRSC surface area covered by plaques found significant main effects of age ($F(2, 34) = 5.44$, $p < .01$; 3-way mixed ANOVA) and genotype ($F(1,34) = 7.86$, $p < .01$), as well as an interaction between the two variables ($F(2,34) = 5.44$, $p < .01$). In the gRSC, area covered was only significantly increased in the 9m group compared to 3m. There was no main effect of sub-region ($F(1,34) = 0.18$, $p = .68$), or interaction between sub-region and age ($F(2,34) = 1.06$, $p = .36$) or genotype ($F(1,34) = 0.18$, $p = .68$). **D,** Analysis of average surface area of plaques in Tg mice showed a significant main effect of age ($F(2,19) = 10.21$, $p < .001$; 2-way mixed ANOVA) but no main effect of sub-region ($F(1,19) = 0.12$, $p = .73$) or interaction of sub-region with age ($F(2,19) = 1.16$, $p = .33$). Boxplots display median (solid line), IQR and range. * *p < .05,* ** *p < .01.*

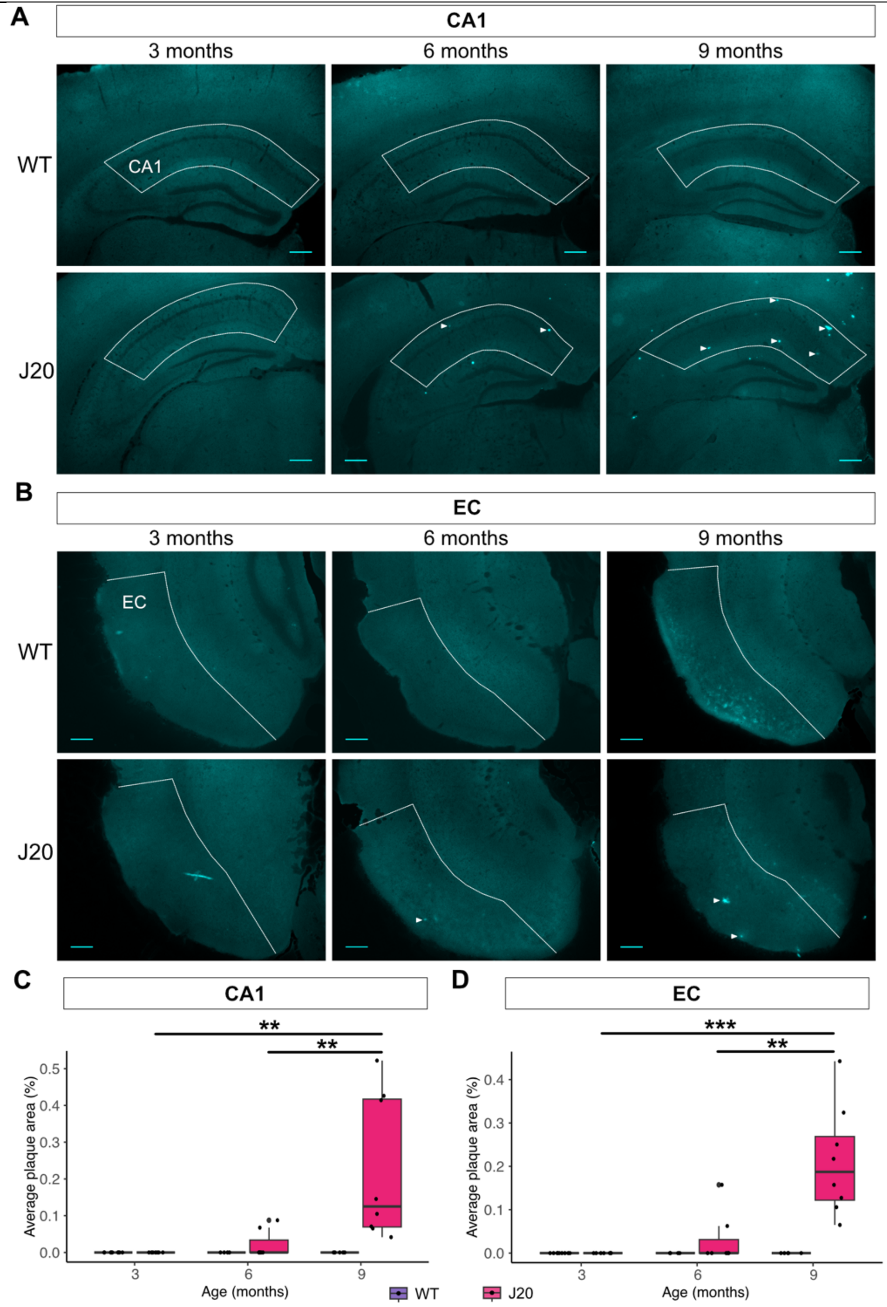


**Supplementary figure 2: amyloid plaque deposition in hippocampus and entorhinal cortex increases with age in J20 mice. A,** representative images of Aβ plaque deposition in the CA1 in WT and J20 mice at 3m, 6m and 9m. Arrows indicate plaques at 6m and 9m in J20 mice. Scale bars: 250 µm. **B,** representative images of Aβ plaque deposition in the lateral EC in WT and J20 mice at 3m, 6m and 9m. Arrows indicate plaques at 6m and 9m in J20 mice. Scale bars: 250 µm **C,** analysis of CA1 surface area covered by plaques found significant main effects of genotype ($F(1,34) = 9.50$, $p < .01$; 2-way ANOVA) and age ($F(2,34) = 8.27$, $p < .01$), as well as a significant interaction between the two ($F(2,34) = 5.97$, $p < .01$). Tg *post hoc* tests found area covered was significantly higher at 9m as compared to both 3m and 6m

(**3m:** 0 (0); **6m:** 0.02 (0.04); **9m:** 0.22 (0.20) %), while 3m and 6m did not differ significantly (*p* = .74). **D,** analysis of EC surface area covered by plaques found significant main effects of genotype ($F(1,34) = 19.20$, $p < .001$; 2-way ANOVA) and age ($F(2,34) = 14.68$, $p < .001$), as well as a significant interaction between genotype and age ($F(2,34) = 10.82$, $p < .001$). *Post hoc* tests showed that plaques covered significantly more EC area in Tg mice at 9m than at 3m and 6m (**3m:** 0 (0); **6m:** 0.03 (0.06); **9m:** 0.21 (0.13) %). There was no significant difference between 3m and 6m (*p* = .55). Boxplots display median (solid line), IQR and range. ** $p < .01$, *** $p < .001$.

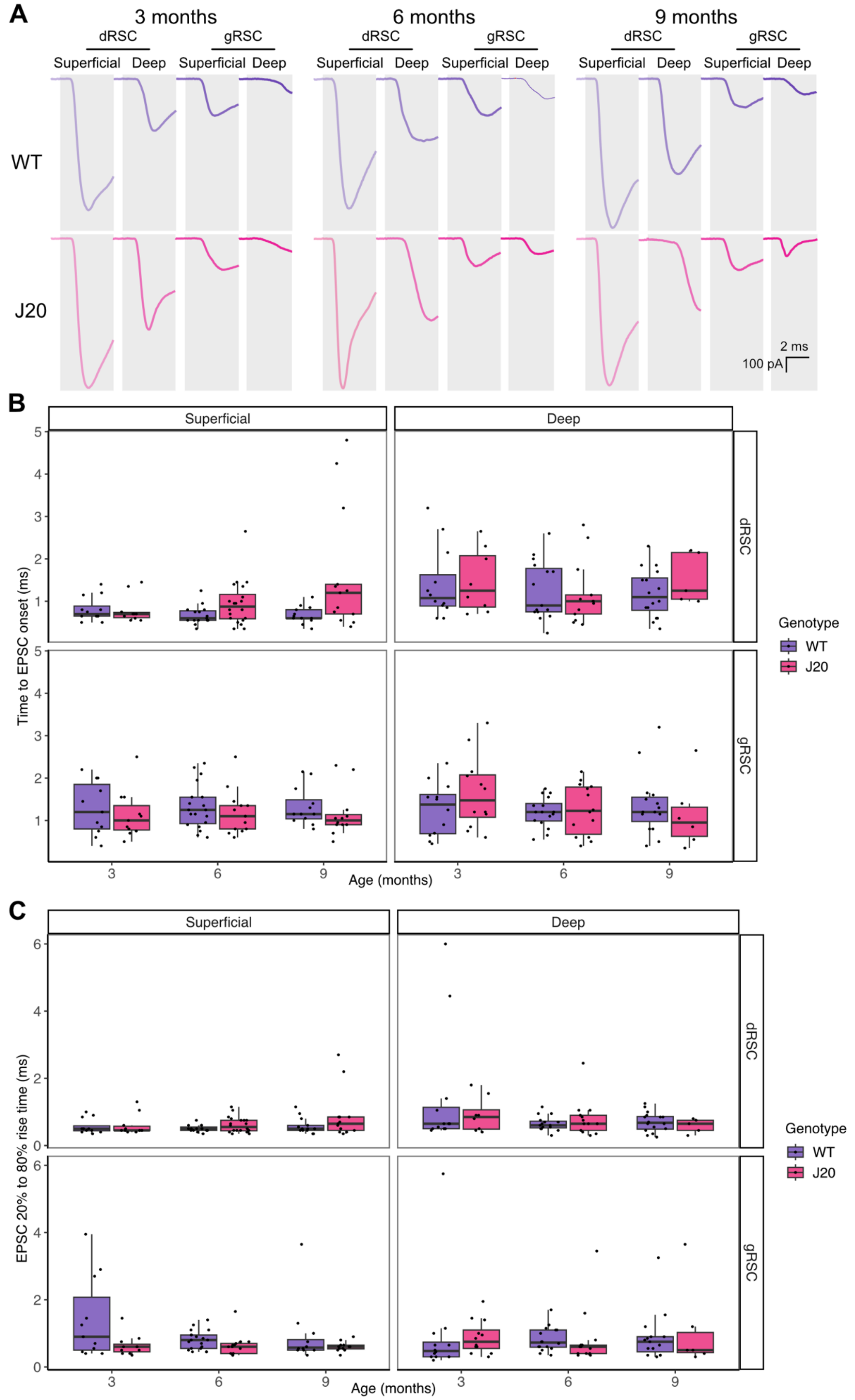


**Supplementary figure 3: no change in ATN-EPSC onset or 20 to 80% rise time in J20 mice or with age. A,** representative voltage-clamp traces ($V_H$ = -70 mV) showing onset of the EPSC generated by the optogenetic stimulation (grey boxes indicate optical stimulation period) of ATN axons. **B,** time to EPSC onset differed between sub-region and layer: dRSC superficial cells appear to show the shortest onset time. Neither age nor genotype significantly explained EPSC onset time variance, and no clear difference in these factors are observed graphically. Boxplots display median (solid line), IQR and range. **C,** EPSC 20-80% rise time did not differ between sub-region, layer, age or genotype. Boxplots display median (solid line), IQR and range.

| | Onset to EPSC | | | | | | | | | | | |
|---|---|---|---|---|---|---|---|---|---|---|---|---|
| | **Model 1** | | | **Model 2** | | | **Model 3** | | | **Model 4** | | |
| *Predictors* | *Estimates* | *t* | *p* | *Estimates* | *t* | *p* | *Estimates* | *t* | *p* | *Estimates* | *t* | *p* |
| Intercept | 0.95 (0.82 – 1.08) | 13.91 | **<0.001** | 0.89 (0.67 – 1.12) | 7.81 | **<0.001** | 0.98 (0.75 – 1.20) | 8.43 | **<0.001** | 1.06 (0.90 – 1.22) | 12.93 | **<0.001** |
| Sub-region: gRSC | 0.22 (0.08 – 0.36) | 3.17 | **0.002** | 0.22 (0.08 – 0.36) | 3.17 | **0.002** | 0.22 (0.08 – 0.36) | 3.17 | **0.002** | 0.22 (0.08 – 0.36) | 3.15 | **0.002** |
| Layer: Deep | 0.28 (0.14 – 0.41) | 3.91 | **<0.001** | 0.28 (0.14 – 0.42) | 3.99 | **<0.001** | 0.28 (0.15 – 0.42) | 4.04 | **<0.001** | 0.28 (0.14 – 0.41) | 3.94 | **<0.001** |
| Age: 6 months | | | | -0.05 (-0.29 – 0.18) | -0.43 | 0.669 | -0.01 (-0.24 – 0.22) | -0.11 | 0.913 | | | |
| Age: 9 months | | | | 0.07 (-0.18 – 0.31) | 0.53 | 0.598 | 0.1 (-0.14 – 0.34) | 0.78 | 0.434 | | | |
| Genotype: Tg | | | | 0.13 (-0.07 – 0.32) | 1.3 | 0.194 | 0.13 (-0.06 – 0.32) | 1.38 | 0.168 | | | |
| Sex: Female | | | | | | | -0.22 (-0.41 – -0.03) | -2.3 | **0.022** | -0.22 (-0.40 – -0.03) | -2.25 | **0.025** |
| **Random Effects** | | | | | | | | | | | | |
| $\sigma^2$ | 0.35 | | | 0.35 | | | 0.35 | | | 0.35 | | |
| $\tau^2_{mouse}$ | 0.08 | | | 0.08 | | | 0.07 | | | 0.07 | | |
| ICC | 0.19 | | | 0.18 | | | 0.16 | | | 0.17 | | |
| N | 68 | | | 68 | | | 68 | | | 68 | | |
| Observations | 305 | | | 305 | | | 305 | | | 305 | | |
| *Marginal $R^2$ / Conditional $R^2$* | 0.070 / 0.248 | | | 0.081 / 0.247 | | | 0.102 / 0.245 | | | 0.091 / 0.244 | | |

**Supplementary table 1: Fixed and random effect results for each mixed model analysing time to onset of EPSC.** For fixed effects (predictors), the table displays effect size, confidence intervals, *t* statistic and significance value. For random effects, the table displays residual variance ($\sigma^2$), mouse variance ($\tau^2$) and the ICC value. Marginal $R^2$ refers to variance explained by fixed effects only, while conditional $R^2$ refers to variance explained by combined fixed and random effects. The best model to explain variability in our data suggested that RSC subdivision, layer and mouse sex all contributed significantly to EPSC onset, but that there was no effect of age or genotype. Model 1 significantly improved upon the null model (random effect of mouse) ($\chi_2(2) = 23.3$, $p < .001$; $\mathbf{AIC_{null}} = 620.5$, $\mathbf{AIC_{M1}} = 601.2$), model 2 did not improve upon model 1 ($\chi_2(3) = 2.7$, $p < .45$; $\mathbf{AIC_{M2}} = 604.5$), model 3 improved upon model 2 ($\chi_2(1) = 5.1$, $p < .05$; $\mathbf{AIC_{M3}} = 601.4$). Finally, model 4 improved upon model 1 ($\chi_2(1) = 4.9$, $p < .05$; $\mathbf{AIC_{M4}} = 598.3$).

| | 20-80% Rise | | | | | | | | |
|---|---|---|---|---|---|---|---|---|---|
| | **Model 1** | | | **Model 2** | | | **Model 3** | | |
| *Predictors* | *Estimates* | *t* | *p* | *Estimates* | *t* | *p* | *Estimates* | *t* | *p* |
| Intercept | 0.66 (0.53 – 0.80) | 9.66 | **<0.001** | 0.98 (0.81 – 1.14) | 11.72 | **<0.001** | 1.02 (0.84 – 1.19) | 11.42 | **<0.00** |
| Sub-region: gRSC | 0.12 (-0.03 – 0.28) | 1.53 | 0.127 | | | | | | |
| Layer: Deep | 0.15 (-0.01 – 0.31) | 1.91 | 0.058 | | | | | | |
| Age: 6 months | | | | -0.23 (-0.42 – -0.04) | -2.35 | **0.019** | -0.21 (-0.40 – -0.02) | -2.14 | **0.03** |
| Age: 9 months | | | | -0.17 (-0.37 – 0.04) | -1.61 | 0.108 | -0.15 (-0.36 – 0.05) | -1.46 | 0.14 |
| Genotype: Tg | | | | -0.09 (-0.25 – 0.07) | -1.14 | 0.253 | -0.09 (-0.25 – 0.07) | -1.13 | 0.26 |
| Sex: Female | | | | | | | -0.1 (-0.26 – 0.06) | -1.24 | 0.21 |
| **Random Effects** | | | | | | | | | |
| $\sigma^2$ | 0.48 | | | 0.49 | | | 0.49 | | |
| $\tau^2_{mouse}$ | 0.01 | | | 0 | | | 0 | | |
| ICC | 0.02 | | | | | | | | |
| N | 68 | | | 68 | | | 68 | | |
| Observations | 305 | | | 305 | | | 305 | | |
| *Marginal $R^2$ / Conditional $R^2$* | 0.020 / 0.041 | | | 0.022 / NA | | | 0.027 / NA | | |

**Supplementary table 2: fixed and random effect results for each mixed model analysing EPSC 20-80% rise time.** For fixed effects (predictors), the table displays effect size, confidence intervals, t statistic and significance value. For random effects, the table displays residual variance (σ2), mouse variance (τ2) and the ICC value. Marginal $R^2$ refers to variance explained by fixed effects only, while conditional $R^2$ refers to variance explained by combined fixed and random effects. No model improved upon the null, suggesting that none of the fixed variables significantly altered 20-80% rise time. Model 1 did not improve on the null model (random effect of mouse) ($\chi_2(2) = 6.0$, $p = .05$; $\mathbf{AIC_{null}} = 660.5$, $\mathbf{AIC_{M1}} = 658.5$), and sequential models did not show significant improvements upon each other ($\chi_2(3) = 6.9$, $p = .08$; $\mathbf{AIC_{M2}} = 659.6$; $\chi_2(1) = 1.5$, $p = .22$; $\mathbf{AIC_{M3}} = 660.0$).

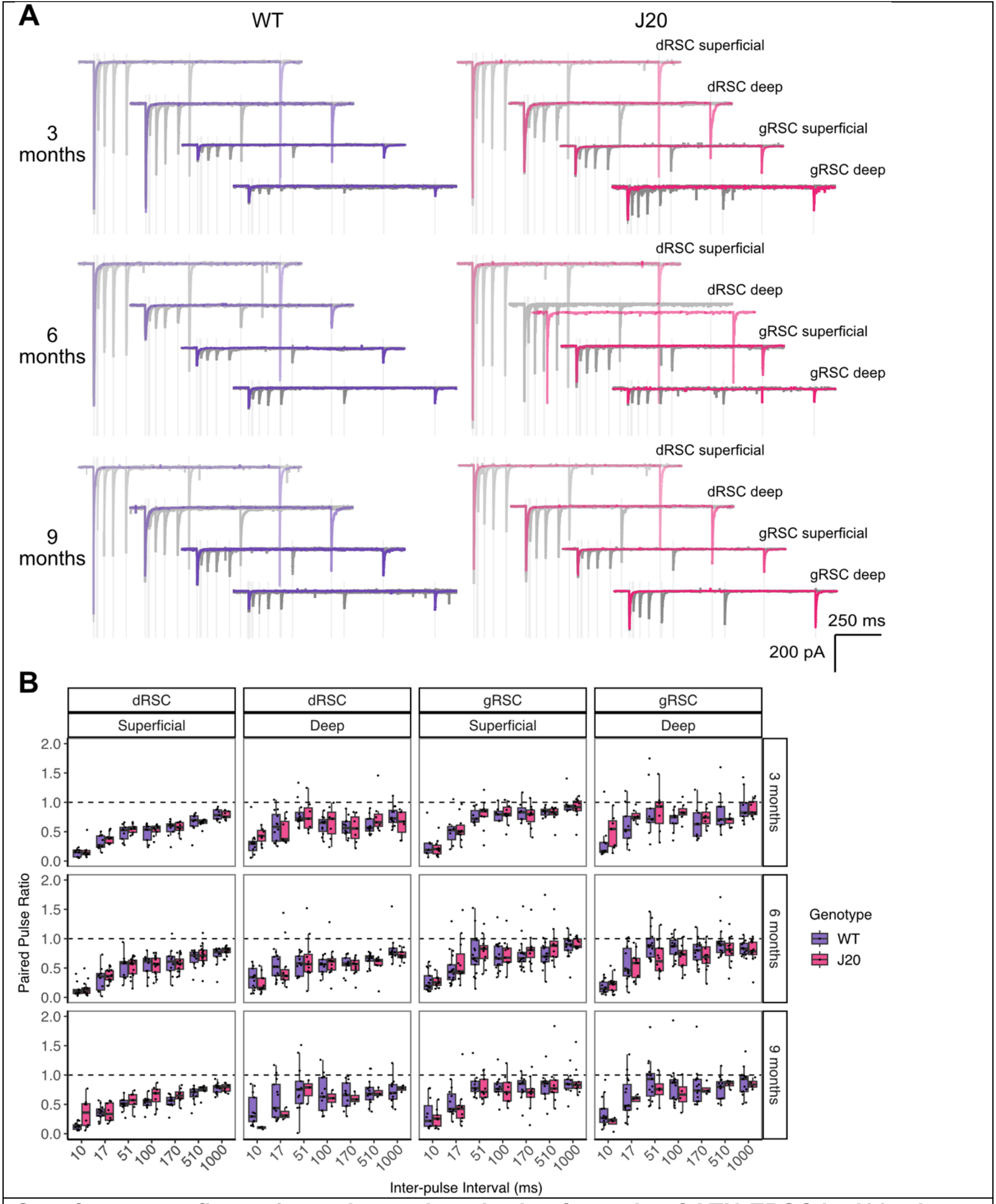


**Supplementary figure 4: no change in paired-pulse ratio of ATN-EPSC in J20 mice or with age. A,** representative voltage clamp traces ($V_H$ = -70 mV) showing EPSCs generated by two optogenetic stimulations separated by increasing intervals. Each plot displays 7 sweeps with intervals of (grey traces) 10 ms, 17 ms, 51 ms, 100 ms, 170 ms, 510 ms and (red trace) 1000 ms. Optical stimulation periods indicated by pale grey boxes. **B,** PPR increased with IPI for all cells, and mean/median PPR remained below 1 (black dashed line). PPR differed between sub-region and layer, and cells in the dRSC superficial layers appear to have a lower PPR which slowly increases. Boxplots display median (solid line), IQR and range.

| | PPR | | | | | | | | |
|---|---|---|---|---|---|---|---|---|---|
| | **Model 1** | | | **Model 2** | | | **Model 3** | | |
| *Predictors* | *Estimates* | *t* | *p* | *Estimates* | *t* | *p* | *Estimates* | *t* | *p* |
| Intercept | 0.18 (0.13 – 0.22) | 7.94 | **<0.001** | 0.19 (0.13 – 0.25) | 6.05 | **<0.001** | 0.18 (0.12 – 0.25) | 5.7 | **<0.001** |
| IPI: 17 ms | 0.23 (0.18 – 0.28) | 9.35 | **<0.001** | 0.23 (0.18 – 0.28) | 9.36 | **<0.001** | 0.23 (0.18 – 0.28) | 9.36 | **<0.001** |
| IPI: 51 ms | 0.44 (0.39 – 0.49) | 17.72 | **<0.001** | 0.44 (0.39 – 0.49) | 17.72 | **<0.001** | 0.44 (0.39 – 0.49) | 17.72 | **<0.001** |
| IPI: 100 ms | 0.41 (0.36 – 0.46) | 16.57 | **<0.001** | 0.41 (0.36 – 0.46) | 16.57 | **<0.001** | 0.41 (0.36 – 0.46) | 16.57 | **<0.001** |
| IPI: 170 ms | 0.4 (0.35 – 0.45) | 16.1 | **<0.001** | 0.4 (0.35 – 0.45) | 16.1 | **<0.001** | 0.4 (0.35 – 0.45) | 16.1 | **<0.001** |
| IPI: 510 ms | 0.48 (0.43 – 0.53) | 19.28 | **<0.001** | 0.48 (0.43 – 0.53) | 19.28 | **<0.001** | 0.48 (0.43 – 0.53) | 19.29 | **<0.001** |
| IPI: 1000 ms | 0.59 (0.54 – 0.64) | 23.71 | **<0.001** | 0.59 (0.54 – 0.64) | 23.72 | **<0.001** | 0.59 (0.54 – 0.64) | 23.72 | **<0.001** |
| Sub-region: gRSC | 0.15 (0.12 – 0.17) | 10.38 | **<0.001** | 0.15 (0.12 – 0.17) | 10.38 | **<0.001** | 0.15 (0.12 – 0.17) | 10.36 | **<0.001** |
| Layer: Deep | 0.04 (0.02 – 0.07) | 3.13 | **0.002** | 0.04 (0.02 – 0.07) | 3.1 | **0.002** | 0.04 (0.02 – 0.07) | 3.09 | **0.002** |
| Age: 6 months | | | | -0.03 (-0.08 – 0.03) | -0.95 | 0.343 | -0.03 (-0.09 – 0.03) | -0.99 | 0.323 |
| Age: 9 months | | | | 0 (-0.06 – 0.06) | 0.07 | 0.947 | 0 (-0.06 – 0.06) | 0.02 | 0.98 |
| Genotype: Tg | | | | -0.01 (-0.05 – 0.04) | -0.23 | 0.816 | -0.01 (-0.05 – 0.04) | -0.23 | 0.815 |
| Sex: Female | | | | | | | 0.01 (-0.04 – 0.05) | 0.36 | 0.723 |
| **Random Effects** | | | | | | | | | |
| $\sigma^2$ | 0.08 | | | 0.08 | | | 0.08 | | |
| $\tau^2_{mouse}$ | 0.01 | | | 0.01 | | | 0.01 | | |
| ICC | 0.07 | | | 0.07 | | | 0.07 | | |
| N | 68 | | | 68 | | | 68 | | |
| Observations | 1743 | | | 1743 | | | 1743 | | |
| *Marginal $R^2$ / Conditional $R^2$* | 0.314 / 0.360 | | | 0.315 / 0.362 | | | 0.315 / 0.363 | | |

**Supplementary table 3: Fixed and random effect results for each mixed model analysing PPR.** For fixed effects (predictors), the table displays effect size, confidence intervals, *t* statistic and significance value. For random effects, the table displays residual variance ($\sigma^2$), mouse variance ($\tau^2$) and the ICC value. Marginal $R^2$ refers to variance explained by fixed effects only, while conditional $R^2$ refers to variance explained by combined fixed and random effects. The paired pulse ratio varied significantly by RSC subdivision and layer, with age and hAPP overexpression having no significant effect.
Model one improved upon the null model (random effect of mouse and within-subject fixed effect of increasing IPI) ($\chi_2(2) = 114.5$, $p < .001$; $\mathbf{AIC_{null}} = 650.0$, $\mathbf{AIC_{M1}} = 539.5$). Sequential models 2 and 3 did not show significant improvement ($\chi_2(3) = 1.6$, $p = .67$; $\mathbf{AIC_{M2}} = 544.0$; $\chi_2(1) = 0.1$, $p = .72$; $\mathbf{AIC_{M3}} = 545.8$).
